# PrecisionTrack: A Platform for Automated Long-Term Social Behavior Analysis in Naturalized Environments

**DOI:** 10.1101/2024.12.26.630112

**Authors:** Vincent Coulombe, Mohamad Sadegh Monfared, Khadijeh Aghel, Quentin Leboulleux, Modesto R. Peralta, FG Blanchet, Benoit Gosselin, Benoit Labonté

## Abstract

Large-scale ethological behavioral studies can provide insights into the neuronal processes underlying complex and social behaviors, potentially opening new avenues for mental health research. However, studying socially interacting animals in naturalistic environments remains technically challenging, as current approaches struggle to simultaneously maintain subject identity, extract behavior, and characterize social interactions over prolonged periods. Here, we present *PrecisionTrack*, an open-source and fully integrated framework designed for real-time multi-animal tracking, behavioral analysis, and social interaction inference in large groups of interacting animals. *PrecisionTrack* achieves high spatiotemporal accuracy in crowded and highly occlusive environments while maintaining robust long-term identity tracking and low-latency processing. To extend behavioral inference beyond pose estimation, we developed the Multi-animal Action Recognition Transformer (*MART*), a transformer-based architecture enabling real-time subject-level action recognition, and *Graph-MART* (*G-MART*), a graph neural network module that infers directed social interactions and interaction partners within groups. In addition, *PrecisionTrack* supports quantitative analysis of evolving social networks across time, enabling investigation of the temporal organization and stability of social dynamics in naturalistic settings. The entire framework is open source and accompanied by standardized workflows and documentation, enabling users to train, evaluate, and deploy custom behavioral analysis pipelines across species and experimental contexts. *PrecisionTrack* provides a scalable platform for quantitative investigation of complex social behaviors at a resolution and duration not accessible with existing methods.

## Introduction

Behavioral studies provide fundamental insights into the neuronal processes that underlie complex constructs such as cognition, emotion, and social interactions. Behavioral analyses are essential not only for understanding the strategies that individuals from different species develop to cope with environmental challenges, but also for identifying how these strategies become maladaptive in models of psychiatric disorders. However, linking these behaviors to underlying neural and social dynamics in complex, naturalistic environments remains a major unresolved challenge.

Traditionally, researchers have relied on manual annotation of predefined ethological features [1– 3]. While effective in controlled settings, this approach becomes impractical in complex social contexts and fails to scale to large cohorts and long-duration experiments. Manual scoring is time-consuming, subject to observer bias, and limits both throughput and reproducibility [4], limitations that become particularly critical when studying social interactions emerging from dynamic interactions between multiple individuals.

The development of computer vision–based artificial intelligence has revolutionized behavioral neuroscience. Tools such as *SLEAP* [5], *DeepLabCut* [6], and others [7,8] use supervised machine learning to automatically track multiple body parts with high spatial and temporal resolution. However, these approaches primarily focus on pose estimation and only partially address the coupling between identity, behavior, and social interactions [5,6], leaving key aspects of multi-individual dynamics unresolved, particularly in dense groups and prolonged recordings.

These limitations emerge across multiple levels of the analysis pipeline. First, current methods often fail to maintain persistent individual identities over time, especially when tracking visually similar animals [9]. Second, many approaches rely on computationally intensive two-stage detection pipelines [7] or are designed for offline analysis [6], limiting their scalability for long-duration experiments. Third, behavioral classification remains largely decoupled from tracking frameworks, requiring separate tools [10,11] that introduce latency and fragment the analysis workflow. Together, these constraints limit the applicability of existing methods to complex, large-scale behavioral studies.

Together, these limitations highlight the inability to simultaneously capture identity, behavior, and social interactions in large groups over extended periods as the main issue with current AI-based approaches for the automatic analysis of complex behaviors. As a result, current approaches fail to capture emergent behavioral structures that arise from interactions within groups.

Here, we introduce *PrecisionTrack*, a unified framework designed to jointly capture identity, behavior, and social interactions in large groups over extended periods. It achieves accuracy comparable to *DeepLabCut* and *SLEAP* while enabling integrated, real-time behavioral and social analysis within a single pipeline. Overall, we present *PrecisionTrack* as a scalable highly integrated architecture enabling long-duration experiments and quantitative analysis of emergent social behavior at a resolution not accessible with existing methods.

## RESULTS

*PrecisionTrack* is an open-source multi-animal pose-tracking and action recognition software (**Figure 1a**) that offers an all-in-one option for behavioral analysis within a unified system (**Figure 1b**). This design enables the transformation of raw, unlabeled behavioral recordings into fully trained, optimized, and deployable analysis pipeline (**Figure 1c**), thereby addressing a central limitation of existing approaches in which tracking, identification, and behavioral inference are typically implemented as separate and hardly coupled tools.

**Figure 1.**
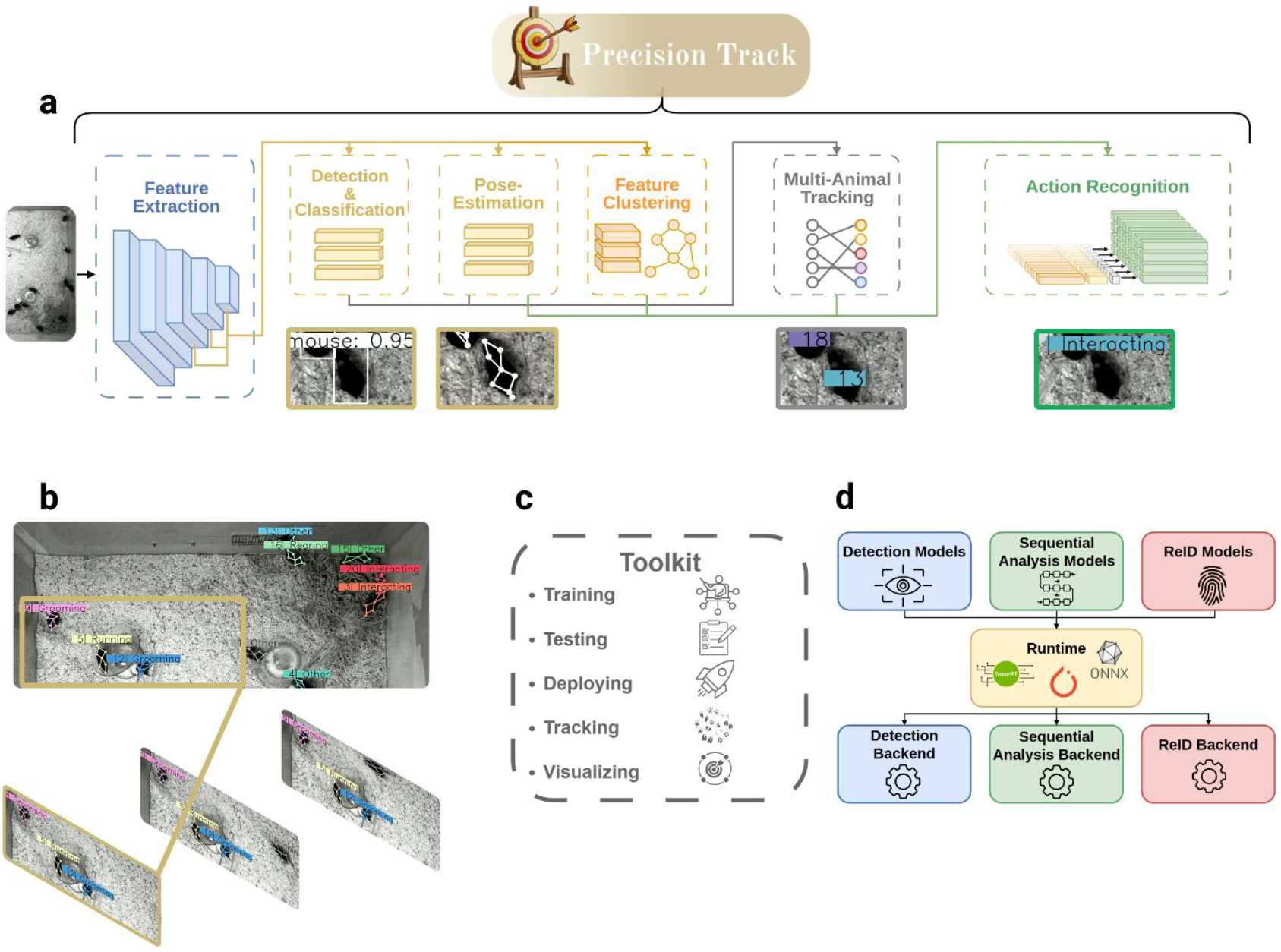
*PrecisionTrack* overview: **a** *PrecisionTrack* executes multiple modules in parallel: feature extraction, subject detection, species classification, pose estimation, multi-animal pose tracking, action recognition, and social behavior classification. **b** *PrecisionTrack* is an online and real-time multi-animal pose-tracker with built-in action recognition and social dynamic analysis. This panel illustrates a qualitative example of its multi-animal pose tracking and action recognition capabilities on the MICE dataset. **c** Overview of the main tools to run or create new predefined, documented pipelines. **d** Illustration of the system’s flexibility. Existing and future algorithms can be incorporated, provided they adhere to the documented interfaces, enabling compatibility with available runtimes and backends.

Within this framework, *PrecisionTrack* provides configurable modules to different species, skeletal definitions, and behavioral repertoires, allowing deployment across a wide range of experimental conditions over prolonged time periods. To ensure interoperability and long-term applicability, the system supports widely adopted runtimes, including *PyTorch* [12], *ONNX* [13], and *TensorRT* [14] (**Figure 1d**), and follow established annotation standards for multi-object tracking. These design choices enable seamless integration with commonly used open-source annotation tools such as *CVAT* [15], *LabelStudio* [16] and *COCOAnnotator* [17].

*PrecisionTrack* is packaged in a reproducible execution environment and hardware-agnostic supporting both CPU and GPU systems with minimal configuration. Combined with comprehensive documentation and standardized workflows, this architecture facilitates rapid adoption and deployment across diverse experimental settings while maintaining the flexibility required to integrate emerging methods and adapt to evolving research needs.

### DATASETS AND BENCHMARKS

Although existing multi-animal datasets have enabled controlled studies of social interactions, they typically involve a limited number of individuals (1–3 animals) in simplified environments, restricting the expression of naturalistic behavioral repertoires [5, 6, 18]. As a result, these datasets do not capture the dense, multi-individual social dynamics that emerge in larger groups and naturalistic settings, and the tracking task they pose is largely saturated by current methods. To address this limitation, we introduce a novel and more challenging dataset, called MICE, designed to better capture the complexity of ethological experimental conditions. More specifically, MICE is composed of recordings of 20 mice freely interacting in a 1m×1m open-field arena enriched with bedding, running wheels, water bottles, feeding trays, tubes, wooden blocks, and nesting material. Each individual is annotated using an eight-keypoint skeleton (snout, left ear, right ear, centroid, left hip, right hip, tail, and tag), enabling detailed pose and identity tracking (**Figure 2a**). Additionally, MICE provide frame-level action annotations (Grooming, Running, Rearing, Interacting, Other) for each visible individual. In total, the dataset contains over one million individually annotated data points, capturing both spatial and temporal dynamics of group behavior at scale. This level of environmental complexity, group size, and annotation density provides a challenging and ecologically relevant benchmark for multi-animal tracking and behavioral analysis (**Figure 2b**). The dataset is publicly available and designed to support standardized evaluation and future methodological developments.

**Figure 2.**
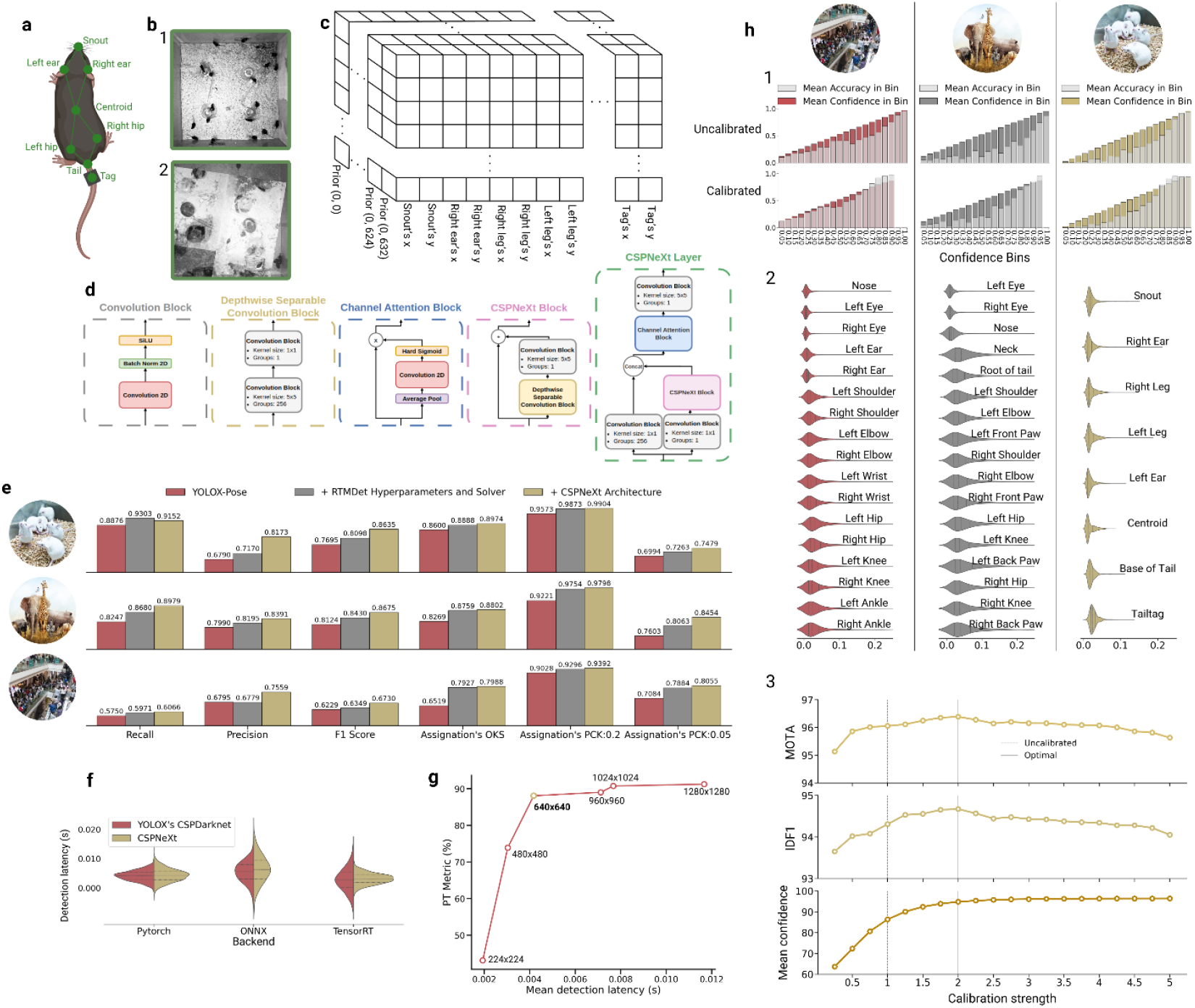
Detection, pose estimation, and calibration: **a** The MICE dataset defines a top-down mouse skeleton with 8 keypoints: snout, left ear, right ear, centroid, left hip, right hip, tail, and tag. **b** (1) Raw training image from the MICE dataset. (2) Augmented training sample from the first training stage, where Mosaic and Mixup augmentations are applied alongside randomly selected geometric and color space transformations. **c** A breakdown of the decoupled keypoint head shows how the prior’s features are processed in parallel, resulting in 8,400 pose predictions across the three levels of the feature pyramid. These predictions are then refined by multiplying them with the prior’s stride and adding the prior’s coordinates. **d** Forward propagation through a *CSPNeXt* layer (green), consisting of *CSPNeXt* blocks (pink), which include a depth-wise convolution block (dimmed yellow), along with a Channel Attention (CA) block (blue) and standard convolution blocks (grey). **e** Performance comparison between the baseline *YOLOX* model (red) and the proposed object detector (dimmed yellow) across three datasets: MICE (top), Animal-Pose (middle), and Microsoft COCO (bottom). Detection quality is evaluated using Recall, Precision, and F1 score, while pose estimation accuracy is assessed with Object Keypoint Similarity (OKS) and Percentage of Correct Keypoints (PCK) at 5% and 20%. Only validation split results are reported. **f** Inference latency comparison. Measured inference delays of *YOLOX*-Pose (red) and the proposed model (dimmed yellow) across different backends (*PyTorch, ONNX, TensorRT*). All measurements were taken at half-precision with a batch size of 1 on an RTX 3090 GPU. It includes preprocessing, forward pass, and post-processing. **g** Detection and pose-estimation score versus inference speed for six models trained at input resolutions ranging from 224 × 224 to 1280 × 1280 pixels. **h** (1) Confidence distribution of the proposed model’s detections on the validation split of the Microsoft COCO dataset (left), the Animal-Pose dataset (middle), and the MICE dataset (right). Detections are grouped into 20 bins, from the least confident (right) to the most confident (left), with the y-axis representing the average confidence within each bin. The pale gray overlay indicates the actual detection accuracy per bin, illustrating the model’s reliability. A well-calibrated model maintains a close to one-to-one accuracy/confidence ratio across a wider upper confidence range. (2) Distribution of keypoint prediction errors for the proposed model across each validation split. Distances are normalized by the subject’s size. (3) Multi-Object Tracking accuracy (MOTA), IDF1 and mean confidence score of the same detection model with more than twenty calibration configurations.

### MULTI-SUBJECT DETECTION, CLASSIFICATION AND POSE-ESTIMATION

We first developed a neural network architecture unifying subject detection, multi-animal tracking, and pose estimation within a single forward pass, using an anchor-free design inspired by YOLO-Pose [21] (**Figure 2c**). By performing detection and pose estimation jointly, this pipeline allows pose estimation to directly benefit from advances in object detection, including strongly augmented training strategies [22-24] (**Figure 2b**). The detection head is based on *YOLOX*, and the backbone adopts a *CSPNeXt* architecture derived from *RTMDet* [25] (**Figure 2d**). *CSPNeXt* leverages large kernel depthwise separable convolutions to expand the effective receptive field without proportional parameter growth, while decoupling channel and spatial mixing improve gradient flow during training. These architectural choices collectively minimize computational overhead while preserving the fine-grained spatial information required for accurate pose estimation in dense, multi-animal scenes.

We evaluated our network’s detection (F1 at 0.5 IoU (**Equation 18**)) and pose-estimation (OKS (**Equation 2**) and PCK) performances against the established *YOLOX-pose* algorithm [21, 22] across three datasets, including MICE, Microsoft COCO [19] and Animal Pose (AP) [20] (**Figure 2e**). Optimizing *YOLOX* training, including replacing SGD [23] with *AdamW* [24], improved performance by 5.49% (**Equation 5**), while substituting *CSPDarknet* with *CSPNeXt* yielded an additional 2.26% gain without affecting inference speed (**Figure 2f**). Noticeably, our inference latency analysis across our supported runtimes: *PyTorch* (4.17ms: 4.28ms), *ONNX* (5.59ms: 6.03ms), and *TensorRT* (3.02ms: 3.14ms) showed no significant differences (**Figure 2f**). We further characterized performance (**Equation 5**) as a function of inference speed across input resolutions ranging from 224×224 to 1280×1280. 640×640 emerged as a Pareto-optimal resolution for detection and pose estimation (**Figure 2g**).

Entropy-based image classifiers are known to exhibit systematic overconfident prediction scores [26]. Our analysis revealed similar overconfidence in our detector’s predictions across all three datasets (**Figure 2h**, Uncalibrated), which could degrade downstream multi-animal pose-tracking relying on confidence scores as proxies for accuracy. To address this, we implemented temperature-scaling calibration [26, 27], reducing confidence–accuracy discrepancies by 16.3%, 23.1%, and 21.8% on MICE, AP, and COCO, respectively (**Figure 2h**). Calibrated confidences improved downstream tracking by reducing both false positives (overconfident low-quality detections) and false negatives (underconfident valid detections), resulting in gains in MOTA (+1.2%) and IDF1 (+0.8%). Please refer to the *robust multi-animal pose-tracking* section for further explanations on these metrics.

Collectively, these results demonstrate that our detection and pose-estimation framework is both accurate and efficient in complex multi-subject environments, providing a reliable and well-calibrated foundation for downstream tracking and behavioral analysis in crowded and highly occlusive experimental conditions.

### FEATURES EXTRACTION AND CLUSTERING

We next asked whether the internal feature representations extracted by our *CSPNeXt* backbone (**Figure 3a**) are sufficiently structured to support downstream tasks such as identity tracking and action recognition. In this context, features refer to representations learned by the network when trained to accurately estimate each subject’s location, pose, and species, rather than hand-crafted descriptors engineered from domain knowledge.

**Figure 3.**
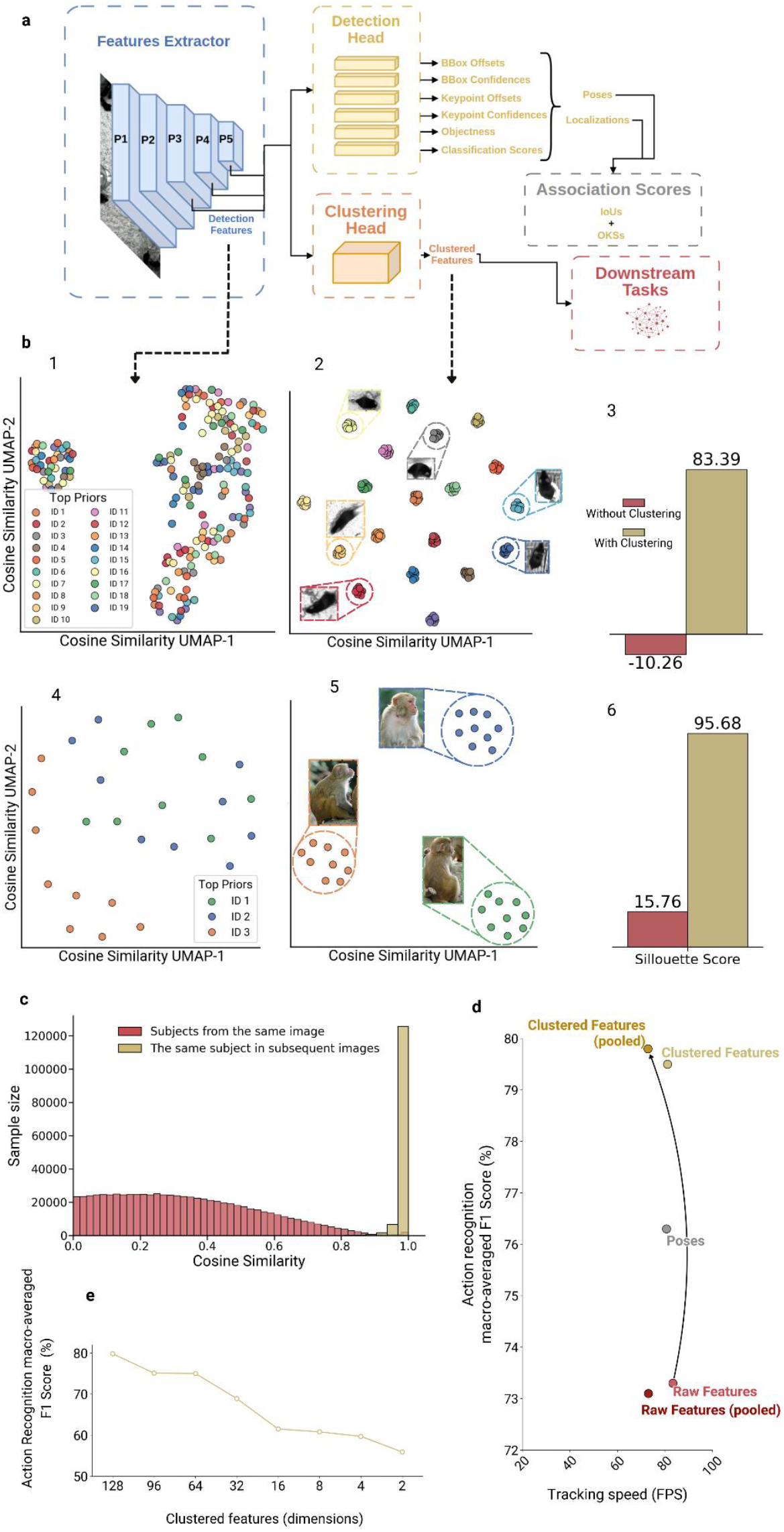
Feature extraction and clustering: **a** Schematic representation of *PrecisionTrack*’s architecture for subject detection, classification, pose estimation, and feature extraction. Raw RGB images are first processed by the feature extractor (blue), producing feature maps. The last three feature maps are selected for further processing. These selected maps are processed by various decoupled heads (yellow) to produce subject poses, bounding boxes, and classification scores. Simultaneously, the same feature maps are processed by the clustering head to extract entity-specific features (orange). **b** Qualitative and quantitative evaluation of the coherence of the detection features and the extracted features. (1, 4) 2D visualization of each detected entity’s top-k detection features from an image in the MICE dataset (top) and the AP dataset (bottom). (2, 5) 2D visualization of each detected entity’s top-k extracted features from an image in the MICE dataset (top) and the AP dataset (bottom). (3, 6) Silhouette scores measuring cluster coherence for detection features (red) and extracted features (dimmed yellow) on the MICE (top) and AP (bottom) datasets. **c** The clustered features belonging to the same individual across two subsequent frames are very similar (dimmed yellow). On the other hand, the clustered features of different individuals within the same frame are dissimilar (red), meaning that subject’s clustered features remain coherent over subsequent frames. **d** Impact of feature clustering on downstream action recognition. Here, “pooled” means that all the features leading to redundant detections are averaged instead of simply selecting the ones linked to the most confident detection. Notably, Pooling clustered features yields measurable accuracy gains, whereas pooling raw, unclustered features degrades performance. **e** Compressing the clustered features correlates negatively with downstream task performance.

High-dimensional data pose a significant challenge in pattern recognition due to their potentially high signal-to-noise ratio [28]. This is particularly true for high-resolution and semantically dense scenes, where multiple visually similar individuals interact and occlude one another [29-31]. However, architectures, such as *CSPNeXt*, combining large effective receptive fields with multi-scale feature aggregation capture efficiently both local details and global contexts [32]. This suggests that our network extracts features rich enough to support context-dependent downstream analyses.

To verify this hypothesis, we evaluated the quality of the extracted features on the task of frame-level action recognition using the MICE dataset. Specifically, we compared the performance of a downstream action recognition network when provided with either the subject’s estimated pose or its features extracted by our *CSPNeXt* backbone. The downstream model (see section *automatic behavioral and social dynamic analysis*) achieved a macro-macro-averaged F1 score of 73.3% when using the extracted features, compared to 76.3% when using the pose alone, meaning that the extracted features could be semantically meaningful enough to complement the subject poses as input to downstream algorithms.

We next investigated the structure of the extracted features in the context of multi-animal tracking, expecting that features associated with redundant detections belonging to the same individual would exhibit high cosine similarity. Surprisingly, we found the opposite. Features leading to highly overlapping detections (ultimately merged into a single detection during post-processing) were often dissimilar, a phenomenon we refer to as feature map incoherence (**Figure 3b**). To quantify this effect, we computed silhouette scores measuring the consistency of representations belonging to the same individual. Initial scores were low (MICE: −10.26%; AnimalPose: 15.76%), confirming poorly clustered embeddings with high intra-subject variability (**Figure 3b**). We therefore hypothesized that improving the coherence of the network’s feature space would improve the performance of downstream algorithms that rely on these representations as input.

To test this without compromising the network’s detection and pose estimation capabilities, we introduced a lightweight convolutional clustering head (**Figure 3a**) trained with a semi-hard triplet loss [33]. Rather than jointly optimizing detection, pose estimation, and clustering, an approach that can degrade primary task performance [35, 36], we adopted a curriculum-learning strategy in which the network is first trained for detection and pose estimation, after which the clustering head is optimized on frozen feature maps. This decoupled training substantially improved feature coherence (**Figure 3b**), with validation silhouette scores reaching 83.39% (MICE) and 95.68% (*AP*), without degrading detection performance. At inference, features belonging to the same cluster are pooled to produce stable identity representations over time.

To evaluate the downstream impact of improved feature coherence we conducted three complementary analyses. First, we compared the temporal and spatial consistency of identity representations and observed that the features, after being clustered by the clustering head, produce stable identity assignments across consecutive frames (**Figure 3c**). Second, we re-evaluated the action recognition task described above using the clustered features as input to the downstream network. The clustered features achieved an action recognition macro-averaged F1 score of 79.8%, with negligible impact on the overall system latency compared to previous inputs (**Figure 3d**). Third, we varied the dimensionality of the clustering head’s output to assess the effect of feature compression on downstream performance. We found that compression negatively correlates with downstream accuracy (**Figure 3e**), suggesting that the richer representational capacity of higher-dimensional embeddings is necessary to preserve the contextual information that benefits action recognition.

### ROBUST MULTI-ANIMAL POSE-TRACKING

A central challenge in the multi-animal tracking field is to maintain persistent identities over time, particularly in dense scenes with frequent occlusions and visually similar individuals. To address this, we tackled the problem from local identity association to long-term identity recovery. First, to enable accurate multi-animal tracking, we adopted a Tracking-By-Detection (TBD) paradigm [37], in which detection is followed by identity association. Among recent association methods [37, 38, 39], *ByteTrack* [40] is a motion-based state-of-the-art. However, it is primarily optimized for structured scenarios in which pedestrians and vehicles move in a unidirectional and relatively steady manner [41] and is therefore not well suited for ethological contexts characterized by frequent path crossings, irregular trajectories, and prolonged occlusions.

To address these limitations, we first augmented *ByteTrack*’s intersection-over-union (IoU)-based association cost with a variant of the object keypoint similarity (OKS) metric (**Equation 13**), allowing identity assignment to leverage both spatial proximity and pose information. As shown in **Figure 4a**, location-only cues are insufficient to maintain reliable identities in highly occlusive scenarios. On the MICE benchmark, augmenting *ByteTrack*’s association with OKS modestly increased MOTA from 95.5% (IoU only) to 95.6% (IoU+OKS) and more substantially improved IDF1 from 88.4% (IoU only) to 90.7% (IoU+OKS).

**Figure 4.**
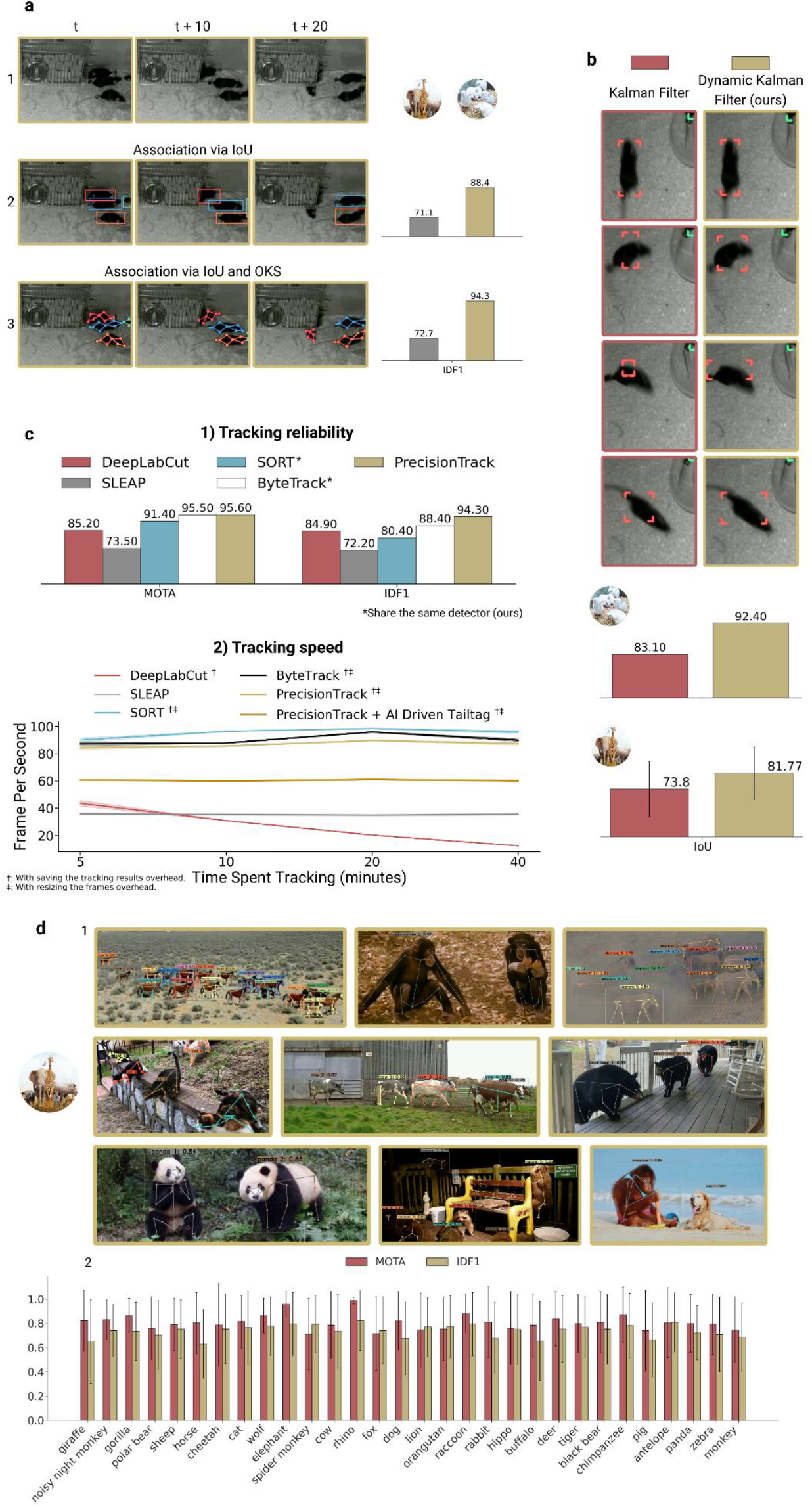
Tracking algorithms and long-duration identification: **a** Representative example illustrating the benefit of OKS-enhanced association. The red mouse is partially occluded by the feeder (1). An algorithm relying solely on IoU would fail to associate the newly visible head (2, t+20) with the red mouse, as there is no overlap with its last seen position (2, t+10). In contrast, an algorithm using both IoU and OKS succeeds, since the ears of the new detection (3, t+20) are close to the red mouse’s last seen ears (3, t+10), enabling the association. **b** Side-by-side comparison between the standard linear Kalman Filter with constant velocity and our Dynamic Kalman Filter. (1) Qualitative comparison when a tracked subject makes a sharp turn followed by rapid acceleration. Corners indicate predicted positions in the next frame, not actual tracking coordinates. (2) Quantitative comparison of predictive performance on the MICE dataset (top) and across all species in the AP dataset (bottom), showing IoU between predicted and actual next frame locations. **c** (1) Multi-Object Tracking Accuracy (MOTA) and IDF1 scores for *PrecisionTrack, DeepLabCut, SLEAP, SORT* and *ByteTrack* on the 20-mice MICE benchmark. All models were trained on the MICE pose-estimation dataset. The selected *DeepLabCut* model is a bottom-up *dlcrnet_ms5* with PAFs enabled; the *SLEAP* model is a bottom-up *EfficientNetB0* with simple max tracks and IoU-based Hungarian matching. *PrecisionTrack, SORT* and *ByteTrack* share the same detector (ours, **Figure 2**). (2) Tracking speed (FPS) comparison of *PrecisionTrack, DeepLabCut, SLEAP, SORT* and *ByteTrack. PrecisionTrack* retains real-time responsiveness even when paired with the data-driven *Tailtag* re identification module (dark yellow). **d** (1) Qualitative assessment of *PrecisionTrack*’s ability to detect, estimate poses, and classify more than 30 species. (2) Quantitative evaluation of *PrecisionTrack*’s Multi-Object Tracking Accuracy (MOTA) and IDF1 when tracking 30 distinct animal species from the APTv2 dataset [43]. Tracking performance is not determined by species, but by viewpoint quality and occlusion level in the recordings.

*ByteTrack* employs a standard linear Kalman filter with a constant-velocity model [42] to predict subject locations across frames. However, animals in ethological environment exhibit erratic movement patterns characterized by sudden changes in both direction and speed. Consequently, we introduced the *Dynamic Kalman Filter* (*DKF*), which incorporates two-dimensional Newtonian dynamics into the original filter. As shown in **Figure 4b**, *DKF* better captures rapid changes in direction and speed typical of animal behavior, improving motion prediction on both MICE (KF: 83.1%; DKF: 92.4%) and AP (KF:73.8%; DKF: 81.8%) datasets and increasing IDF1 from 90.7% (IoU+OKS only) to 94.3% (IoU+OKS+DKF) on MICE.

Finally, we benchmarked *PrecisionTrack* against established multi-animal pose tracking (*DeepLabCut* [6], *SLEAP* [5]) and MOT baselines (*SORT* [37], *ByteTrack* [40]) on our MICE dataset under identical training conditions. We evaluated the method’s tracking accuracy the official MOT challenge [41] metrics. More specifically, the Multi-Object Tracking Accuracy (MOTA) (**Equation 21**), which measures the system’s ability to accurately detect subjects in videos without switching them, and the IDF1 (**Equation 22**), which calculates an F1 score based on the tracked subject’s predicted identities, metrics were used. We also evaluated the method’s end-to-end tracking throughput (FPS) on videos of up to 20 animals over 5–40 minutes. *PrecisionTrack* achieved performance on par with or exceeding existing methods, while maintaining real-time processing capabilities. **Figure 4c** further details comparative performance across methods, showing MOTA (*PrecisionTrack*: 95.6%; *DLC*: 85.2%; *SLEAP*: 73.5%; *SORT*: 91.4%; *ByteTrack*: 95.5%) and IDF1 (*PrecisionTrack*: 94.3%; *DLC*: 84.9%; *SLEAP*: 72.2%; *SORT*: 80.4%; *ByteTrack*: 88.4%) on our MICE benchmark. *PrecisionTrack* preserves low processing latency as experiment duration increases. For instance, processing 40 minutes of footage pre-recorded at 25 FPS required 12.5 minutes with *PrecisionTrack*, compared to 29 minutes with *SLEAP* and 142 minutes with *DeepLabCut*. Comparable evaluations for *SORT* and *ByteTrack* yielded processing times of 11.7 and 12.1 minutes, respectively (**Figure 4c**). In addition, to assess the generalization capacity of our framework, we evaluated tracking performance on the Animal Pose tracking (APTV2) dataset [43], a video-based benchmark comprising 2,749 video clips from 30 species, specifically designed to evaluate pose estimation and tracking jointly across temporal sequences. We observed stable tracking performance across species (**Figure 4d**), indicating that *PrecisionTrack* is species agnostic and supports multi-species applications.

Overall, these results indicate that *PrecisionTrack* maintains high tracking accuracy in crowded, naturalistic environments while enabling real-time analysis of long-duration experiments, achieving competitive tracking performance and addressing a key issue in multi-animal tracking by enabling reliable identity tracking over time in dense social settings.

### SUBJECT RE-IDENTIFICATION

Previous sections established that *PrecisionTrack* achieves high accuracy and robustness in crowded, enriched environments. Nevertheless, prolonged occlusions remain a fundamental limitation for multi-animal tracking, leading to identity switches and fragmented trajectories. Such events occur when animals enter nests, dig into bedding, or move within enrichments and degrades long-term IDF1 performance. To mitigate this issue, we first developed a trajectory stitching algorithm that estimates a subject’s likely reappearance region based on its recent motion. Specifically, dynamic search zones are defined around each subject’s last known position and expand over time according to its last known 2D motion dynamics (**Figure 5a**). This approach preserves identity continuity when a single subject reappears (**Figure 5a**). However, in scenarios where multiple subjects reappear within overlapping search zones, stitching alone becomes insufficient (**Figure 5a**), with re-identification accuracy decreasing to 40.8% when six mice re-emerge in the same area.

**Figure 5.**
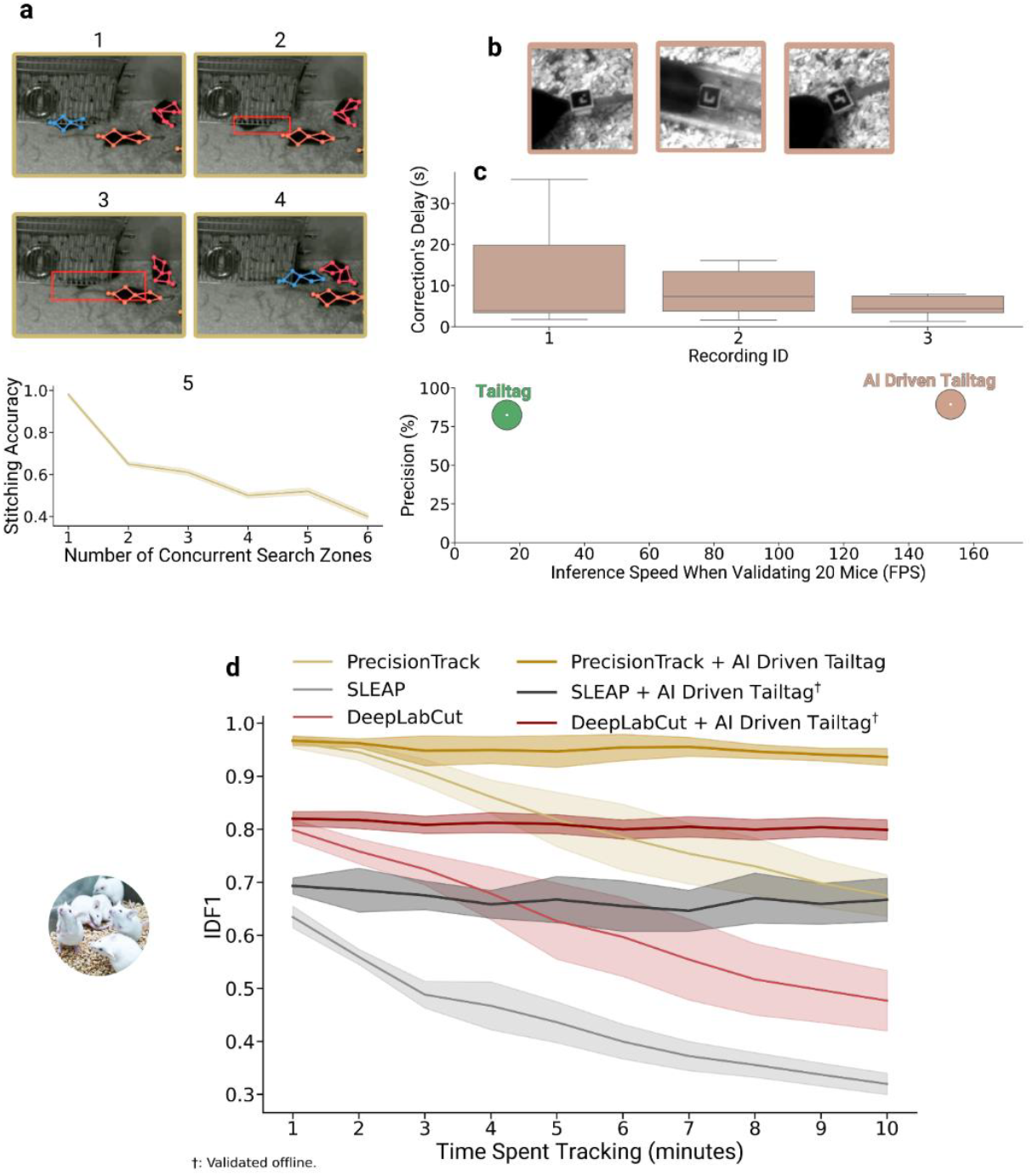
Trajectory stitching and subject re-identification: **a** This algorithm leverages lost subjects (blue) last seen dynamics and location (1), then recovers its trajectory by incrementally expanding the assigned search zones (2, 3) and selecting the best fit once a new subject is found (4). The algorithm’s effectiveness rapidly degrades when multiple subjects are lost at the same location, leading to many overlapping search zones (5). **b** Visual representation of the improved marker detection context. The *Tailtag* system now searches for markers only in regions of interest, rather than across the full frame. **c** (1) Distribution of correction delays across three 10-minute recordings. (2) Reading precision (y-axis) and speed (x-axis) comparison between the improved detection context (dimmed pink) and the original *Tailtag* system (green). By limiting marker searches to relevant regions, the system significantly reduces computational load and improves overall speed. **d** Evolution of the IDF1 tracking metric over 10-minute recordings for *PrecisionTrack* (dimmed yellow), *DeepLabCut* (red), and *SLEAP* (grey). We also evaluated each method when paired with the data-driven *Tailtag* system. All three methods are able to maintain their performance levels when paired with the *Tailtag* system.

Under these conditions, identity recovery becomes fundamentally ambiguous using only motion cues, motivating a more robust re-identification strategy. When subjects are visually indistinguishable, purely markerless re-identification is inherently unreliable. We therefore employed visual tagging as a principled solution and integrated the *Tailtag* system [44], a lightweight marker designed for long-term use without behavioral interference. *Tailtags* are visible under both daylight and infrared conditions and do not require specialized imaging hardware [44]. We integrated *Tailtag* into *PrecisionTrack* and introduced two optimizations. First, we restricted tag-based identification from the full frame to local regions around the predicted tag positions (**Figure 5b**), which improved inference speed without compromising precision (**Figure 5c**). Second, we replaced the original *Tailtag* reassignment procedure with a data-driven tag reading strategy that accumulates evidence over time and updates identities only once a confidence threshold is reached.

We evaluated re-identification performance using our MICE longitudinal benchmark, comprising over one million manually curated mouse localizations. Without the data-driven *Tailtag* re-ID module, *PrecisionTrack, SLEAP* and *DeepLabCut* show rapid decreases in their ability to maintain correct identification, as measured by IDF1 (**Figure 5d**). In contrast, when paired with our data-driven *Tailtag* system, all three were able to sustain their initial identification capabilities (*PrecisionTrack*: mean IDF1 (mIDF1) after 1 minute; 95.7%, mIDF1 after 10 minutes; 93.3%: *SLEAP*: mIDF1 after 1 minute; 69.7%, mIDF1 after 10 minutes; 66.6%: *DeepLabCut*: mIDF1 after 1 minute; 82.83%, mIDF1 after 10 minutes; 79.89%; **Figure 5d**). Furthermore, our analysis shows that our data-driven *Tailtag* system corrects most tracking errors within a few seconds, with more than 90% of errors corrected within 10 seconds and the remaining within 30 seconds (**Figure 5c**), with outliers often corresponding to cases where mice hid immediately after a tracking error.

Globally, our results show that implementing a two-stage re-identification process, combining our stitching algorithm with our data-driven *Tailtag* system, effectively overcomes one of the most important limitations of modern vision-based trackers, namely, their inability to accurately track near-identical subjects in crowded settings over prolonged periods.

### AUTOMATIC BEHAVIORAL AND SOCIAL DYNAMIC ANALYSIS

Accurately identifying individual actions and social behaviors is essential for understanding how animals interact, adapt, and organize within their environments. However, current multi-animal tracking frameworks lack integrated support for action inference. Consequently, researchers rely on standalone tools [10,11] for post hoc behavioral classification, resulting in fragmented pipelines, increased computational cost, offline processing constraints, and manual intervention. To address this limitation, we integrated a dedicated action recognition module into *PrecisionTrack*, termed *MART* (Multi-animal Action Recognition Transformer). Operating in parallel via the *ParallelTracker* API, *MART* enables subject-level inference of actions and social behaviors without compromising *PrecisionTrack*’s real-time performance, thereby extending the framework from tracking to fully integrated behavioral analysis.

*MART* is a transformer decoder (**Figure 6a**) structurally similar to *GPT-2* [45], widely used for sequence modeling in natural language processing [46]. We formulate action recognition as a temporal sequence classification problem over subject level representations, enabling *MART* to integrate temporal context for behavior inference. The temporal extent of this integration is controlled by the context window, which determines how far back in time the model attends. Our analysis reveals that a context window of 30 frames (~1 s) provides optimal performance for mouse action prediction. Allowing larger temporal context substantially improves accuracy relative to single-frame inference (Context window 1: 79.5%; 15: 84.1%; 30: 84.3%; 60: 83.5%; **Figure 6b**). We next assessed whether enriching per-timestep representations improves performance.

**Figure 6.**
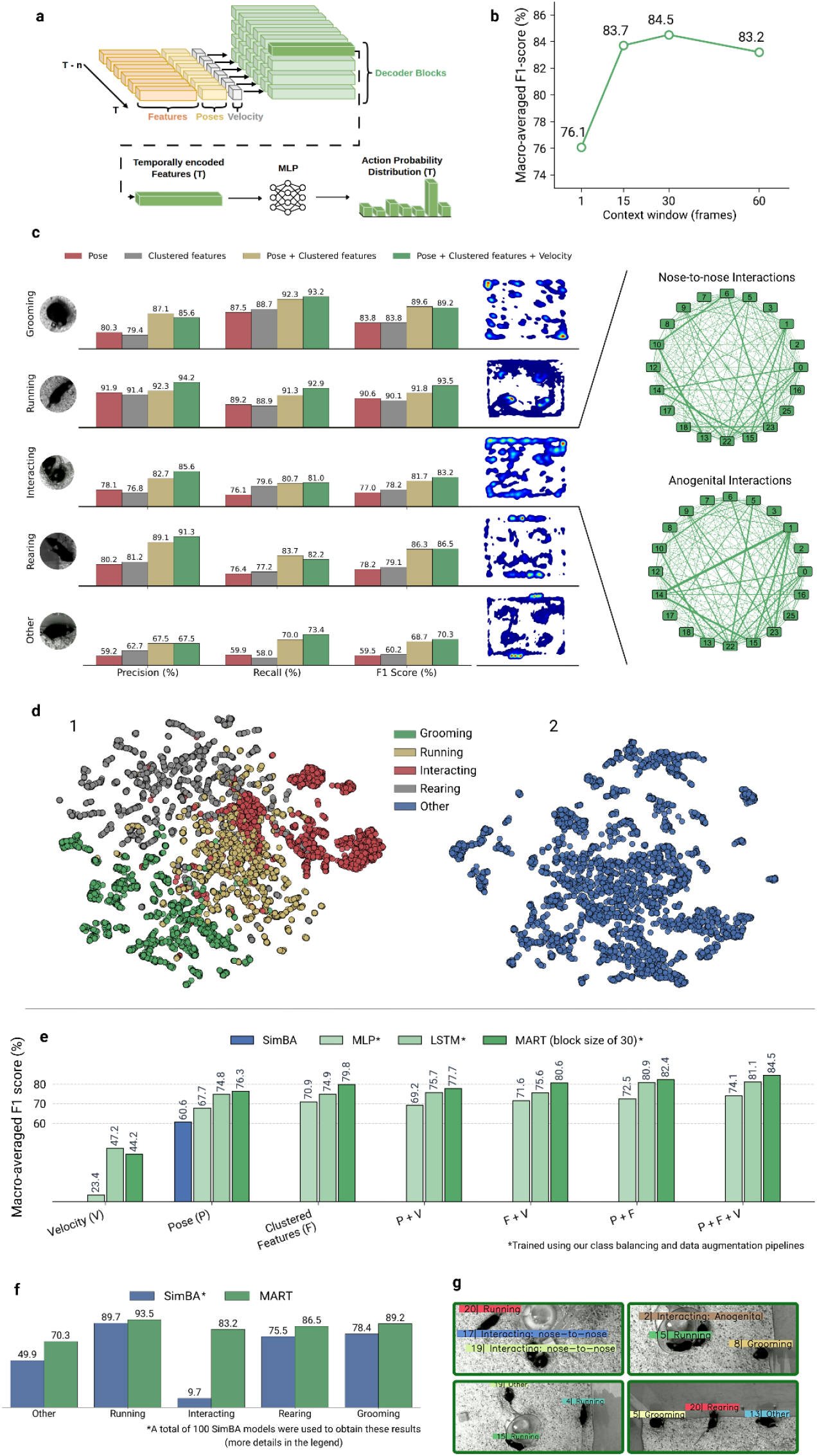
*MART* architecture, ablations, and behavioral analysis: **a** Overall architecture of the Multi-animal Action Recognition Transformer (*MART*). The context window (containing n frames) spans the z-axis, the representations lie along the x-axis, and the four decoder blocks are shown on the y-axis. **b** Context window ablation study, showing the macro-averaged F1 score of each tested context window (or block size). **c** Representation ablation study for all labeled actions and the “Other” action, which serves as the negative class. Recall, Precision, and F1 scores are shown for each representation. Action-specific heatmaps indicate spatial hotspots within the vivarium. Using *PrecisionTrack* poses, we refined some interactions to distinguish between nose-to-nose and anogenital interactions, enabling the construction of social networks depending on the nature of the social interactions. **d** The final decoder block of *MART* can cluster both labeled actions (1) and unlabeled actions (2). **e** Model and input ablation study comparing three architectures (*MLP, LSTM*, and *MART*) across all possible pose–velocity–feature input combinations. The x-axis enumerates all the possible input combinations containing clustered features (F), pose (P) and/or velocity (V). *SimBA* ensemble performance is shown alongside models trained using poses only. **f** Action recognition F1 scores (%) across the five labeled action classes present in the MICE dataset for our *SimBA* models ensemble (blue) as well as *MART* (green). The ensemble was required because *SimBA* performs binary classification (one class vs all) on a per-frame (not per-subject) basis. Therefore, we trained one *SimBA* model per class (5) and selected the highest-confidence prediction for each subject (20) during evaluation, which gives a total of 100 predictions per image. **g** Qualitative example of *MART* detecting individual behaviors (grooming, running, rearing) and social behaviors (nose-to-nose and anogenital interactions).

Using poses (**Figure 6c**, red) or learned clustered features (**Figure 6c**, grey) alone yields comparable results (clustered features (macro-averaged): 79.8%; poses (macro-averaged): 76.3%), whereas combining both produces a clear gain (**Figure 6c**, dimmed yellow (macro-averaged); 82.4%). Incorporating velocity further improves performance (**Figure 6c**, green (macro-averaged); 84.5%), especially for dynamic behaviors such as running and interacting (**Figure 6c**, green; running: 93.5%; interacting: 83.2%). Although not explicitly optimized for clustering, *MART* organizes both labeled classes and a subset of unlabeled “Other” behaviors into coherent groups (**Figure 6d**). **Figure 6g** provides representative examples of action recognition in socially interacting mice within a naturalistic environment.

To rigorously evaluate the performance of *MART*, we benchmarked it against standard architectures commonly used for behavior classification: a multilayer perceptron (MLP) and a long short-term memory network (LSTM) [47]. The MLP provides a feedforward baseline operating on single-frame representations without temporal integration [48], whereas *LSTM* models sequential dependencies through recurrent dynamics and serves as a widely used approach for temporal behavior modeling [49]. All three models were trained under identical conditions and share a similar parameters count (*MART*: 4.4M; MLP: 4.5M; LSTM: 4.4M). **Figure 6e** summarizes the resulting macro-averaged F1 scores across input configurations (velocities, poses, features, and their combinations). *MART* outperforms both alternatives, highlighting the benefit of integrating long-range temporal context through self-attention for behavioral classification.

To further position *MART* relative to existing approaches, we compared its performances against *SimBA* [10], a widely used behavioral classification framework, on our MICE action recognition dataset. Because *SimBA* supports only binary classification and generates a single prediction per frame rather than per subject, we constructed an ensemble of 100 independent classifiers (5 behaviors × 20 subjects). During inference, each subject’s behavior was assigned based on the classifier with the highest confidence score. All *SimBA* models and *MART* were trained and evaluated on identical data. As shown in **Figure 6e**, *MART* achieves higher performance using pose input alone (macro-averaged F1 score: 76.3% vs. 60.6%), while natively providing subject-level, multi-class predictions within a single unified architecture. This eliminates the need for external post-processing pipelines and substantially simplifies behavioral analysis at scale.

A per-class analysis reveals that *SimBA*’s lower overall performance is largely driven by its inability to classify social interactions (Interacting: 9.7%), while its performance on non-social behaviors remains competitive (Running: 89.7%; Grooming: 78.4%; Rearing: 75.5%; Other: 49.9%). This discrepancy stems from a practical constraint: *SimBA*’s cross-animal features, which encode pairwise spatial relationships between individuals, could not be enabled in our experimental setup as its model exceeds the available memory of our training infrastructure (128 GB RAM) given the number of subjects (20) in the MICE dataset. When restricting the evaluation to non-social classes only, the performance gap narrows considerably (*SimBA*: 73.4%; *MART* (pose-only): 76.3%), indicating that *SimBA* remains a strong baseline. However, this comparison underscores a key architectural advantage of *MART*: by operating on temporal sequences of per-subject representations within a unified framework, *MART* captures the contextual information necessary for social behavior classification without requiring explicit pairwise feature engineering or prohibitive memory costs. A detailed per-class comparison between *SimBA* and *MART* across all behavioral categories is provided in **Figure 6f**.

When combined with *MART, PrecisionTrack* enables the spatial and social contextualization of behavior, providing direct insight into how animals organize their activity within complex environments. Mapping action labels onto the vivarium revealed structured behavioral patterns, with rearing occurring predominantly near elevated water bottles and along the perimeter, social interactions clustering around nesting areas, running concentrated on wheels, and grooming primarily observed in isolated regions (**Figure 6c**). Beyond spatial distributions, integrating *MART*’s action labels with *PrecisionTrack*’s pose estimates allowed refinement of social interactions, distinguishing nose-to-nose from anogenital contacts and enabling the construction of interaction-specific social networks (**Figure 6c**). These analyses illustrate how *PrecisionTrack* extends beyond tracking to support quantitative investigation of spatial organization, interaction structure, and emergent social dynamics.

While *MART* enables the identification of actions across individuals, it does not capture the directional nature of social interactions. In ethological research, distinguishing the initiator of an interaction from its recipient is critical, as these roles often reflect distinct underlying neural and behavioral processes [50]. To address this limitation, we developed a spatial encoder inspired by the GraphFormer framework [51], termed *Graph-MART* (*G-MART*). *G-MART* operates on the temporally encoded action features produced by *MART* (**Figure 7a**) and refines them by constructing a directed interaction graph, where nodes represent subjects and edge weights are defined using keypoint-based spatial priors (**Figure 7b**). Through attention-based message passing, node representations are updated based on their relational context, enabling the model to jointly infer both the presence of an interaction (via node classification) and its directionality (via edge classification) using a multi-task focal loss [52].

**Figure 7.**
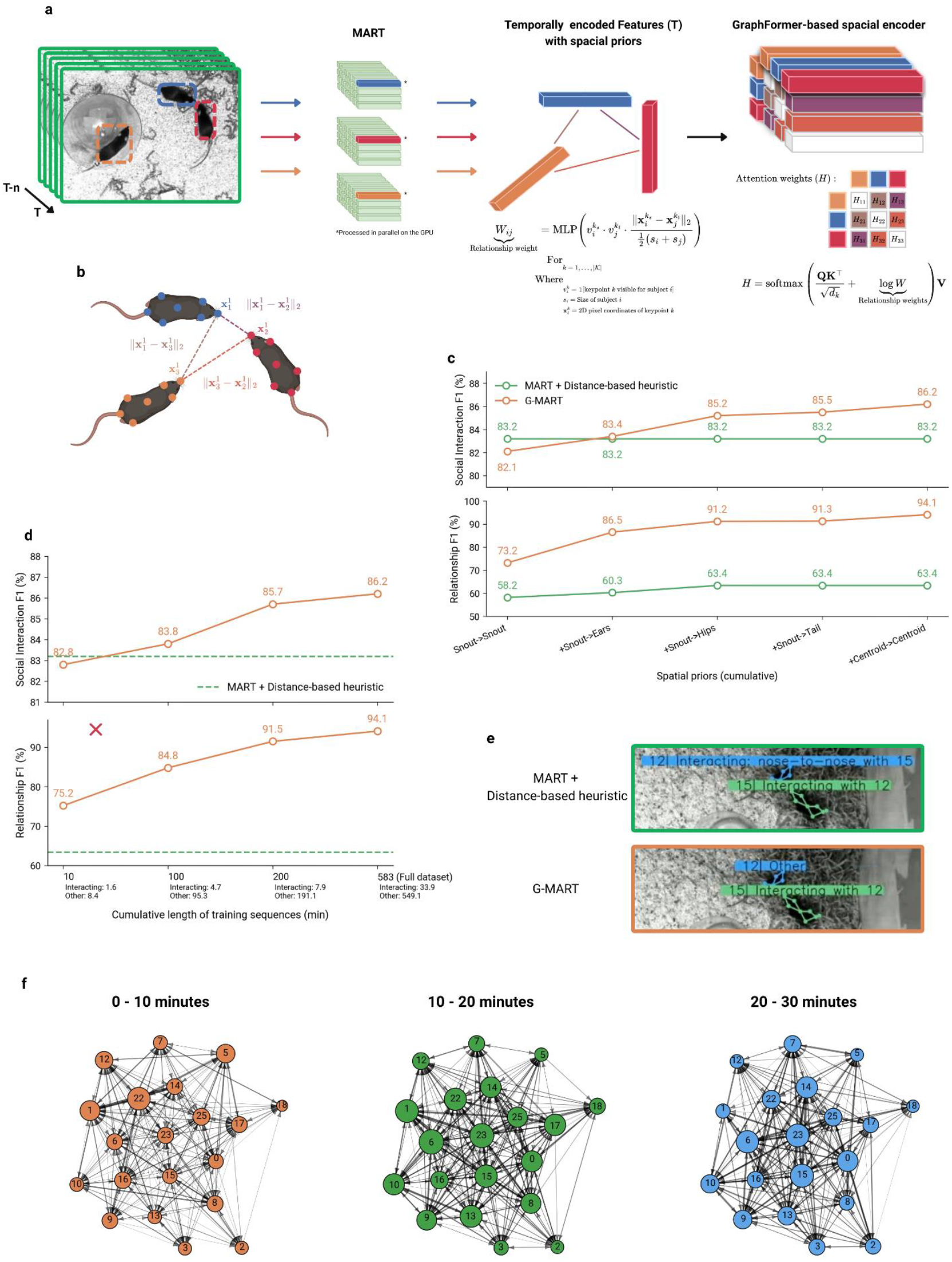
Graph-*MART* and social interaction analysis: **a** *G-MART* architecture. Per-subject poses, features, and velocities are first temporally encoded by *MART*’s decoder, then assembled into a spatial relationship graph where the nodes are the representations of the decoded subject and the edges the relationship weights derived from user-defined keypoint-based spatial priors. This graph is then encoded using a *GraphFormer*-inspired [51] model. **b** An example of snout-to-snout only keypoint spatial priors. *G-MART* builds its spatial relationship graph using relationship weights derived from these user-defined priors. **c** User-defined keypoint-based spatial prior ablation study. Five *G-MART* models were trained with progressively richer sets of spatial priors. The best social interaction detection and relationship prediction performance was achieved using the following spatial priors: snout-to-snout, snout-to-ear, snout-to-hip, snout-to-tail, and centroid-to-centroid distances. **d** Dataset subsampling study. Four *G-MART* models were trained on four different subsamples of the MICE dataset (10, 100, 200 and 583 minutes; 33.9 minutes of interactions total). Extrapolation indicates that *G-MART* surpasses *MART*’s interaction detection performance with approximately 10–20 minutes of labeled data while improving upon the distance-based heuristic in term of relationship prediction. **e** A qualitative example of a scenario where *G-MART* refines and improves upon *MART*’s social action detection capabilities. Here, *MART* (top) incorrectly predicts that subject 12 is interacting with subject 15, triggering the association heuristic. In contrast, *G-MART* (bottom) correctly predicts that only subject 15 is interacting. **f** Continuous time movement model [53] result among all mice using *PrecisionTrack* data over three consecutive 10 minutes blocks. Each node is associated to an individual and the size of each node is representative of the number of interactions of an individual (instigated and received). The direction of an arrow represents the direction of the interaction from giver to receiver, and the width of the arrow characterizes the strength of the interaction (see Method section for more details). The absence of an arrow means that a mouse never instigated an interaction.

Empirically, *G-MART* improved *MART*’s social behaviors detection performance by 3 percentage points (*MART*: 83.2; *G-MART*: 86.2) when *G-MART*’s network’s spatial priors are the following: snout–snout, snout–ears, snout–hips, snout–tail, and centroid–centroid. These user-defined priors reflect ethologically relevant body-part relationships through which mice typically initiate and sustain social contact. **Figure 7c (top)** presents an ablation across spatial prior combinations. Beyond action classification, we also evaluated *G-MART*’s ability to correctly identify relationships (determine which individuals are engaged with a given subject performing a social behavior). Since the only social interaction class in the MICE dataset is “Interacting,” a natural baseline consists in assigning interaction partners based on the spatial priors themselves, referred to as the *MART + distance-based heuristic*. As shown in **Figure 7c (Bottom)**, *G-MART* systematically outperforms this baseline across all spatial prior configurations, demonstrating that learned relational reasoning over directed graphs captures interaction structure more faithfully than geometric heuristics alone.

Finally, we assessed the amount of labeled data required for *G-MART*’s learned representations to surpass the geometric baseline. By progressively subsampling the MICE training set, we found that approximately 20 minutes of annotated footage is sufficient for *G-MART* to outperform the baseline on social behavior classification (**Figure 7d, e**). Notably, *G-MART* improves relationship detection even at the smallest data fractions, likely due to the strong inductive bias provided by its spatial priors. Moreover, performance on both tasks continues to increase with additional training data without signs of saturation, suggesting that larger annotated datasets would yield further improvements.

Leveraging *G-MART*, we obtain refined and directed interaction maps, enabling quantitative analysis of social structure. For example, mouse #1 exhibited frequent anogenital interactions with mice #14 and #22, while interactions with mouse #14 were predominantly nose-to-nose (**Figure 6c**). These results demonstrate that *G-MART* extends behavioral analysis from action recognition to interaction-level inference, capturing both the presence and directionality of social behaviors. Together, *MART* and *G-MART* complete a unified pipeline that links identity tracking, action recognition, and directed interaction analysis. This integrated framework enables scalable and quantitative investigation of complex social dynamics in naturalistic environments, providing a foundation for high-throughput behavioral studies.

Finally, we used *PrecisionTrack* to evaluate the temporal organization of social networks within a colony of mice evolving in a naturalistic vivarium environment. More specifically, we investigated how interaction structure, directionality, and social centrality evolved across three successive 10-minute time periods, enabling quantitative assessment of the stability and dynamics of group social organization over time. Using a continuous-time movement model [53] to quantify directed social interactions, we found that social dynamics within the group were highly variable over time (**Figure 7f**). Both network structure and interaction strength changed substantially across successive 10-minute time blocks. For example, during the first 10-minute interval, mouse 15 occupied a relatively peripheral position within the network, exhibiting few and weak incoming or outgoing interactions, as reflected by the sparse and thin edges connected to node 15 in both directed graphs (**Figure 7f**). However, during the second and third intervals, mouse 15 progressively became more central within the social network, engaging in stronger and more frequent interactions. Conversely, mouse 22 showed the opposite trend, with progressively weaker and fewer interactions over time.

This approach also captures the temporal evolution of reciprocal interactions within the group. In the first 10-minute interval, reciprocal interactions were relatively sparse compared to the latter two intervals, as illustrated by the lower density of bidirectional edges between the corresponding directed graphs (**Figure 7f**). Beyond global network organization, the continuous-time movement model also enables fine-scale analysis of interactions between specific individuals. For instance, interactions between mice 1 and 14 were initially strong and reciprocal during the first interval, became weaker and predominantly unidirectional during the second interval, with mouse 14 primarily initiating the interaction, and were no longer detected during the final interval.

Together, these analyses demonstrate that combining *PrecisionTrack* with continuous-time movement models enables quantitative investigation of the temporal organization, stability, and directionality of social networks. This framework provides new opportunities to study how individuals dynamically shape group structure and social organization over time in naturalistic environments.

## DISCUSSION

The ethological study of behavior across species has advanced our understanding of behavioral repertoires and social strategies that support adaptation to environmental challenges. Recent multi-animal tracking methods enable a shift from manual annotation to automated analyses, yet important limitations still restrict their broader use. In particular, algorithms struggle to re-identify physically similar individuals [5-8], especially under prolonged occlusions, limiting applicability to simplified settings. Although markerless approaches perform well with visually distinct subjects [54], their performance degrades for near-identical animals. In addition, many methods prioritize accuracy over scalability, relying on two-stage or offline pipelines that become impractical for long or dense recordings [6, 7]. Finally, behavioral inference is typically decoupled from tracking, requiring post hoc tools [10, 11] and resulting in fragmented workflows. Together, these limitations highlight the lack of an integrated solution capable of jointly capturing identity, behavior, and social interactions over extended periods. To address these challenges, we developed *PrecisionTrack*, a unified framework for multi-animal tracking and behavioral analysis in naturalistic settings. By integrating identity preservation, real-time processing, and behavior classification within a single architecture, *PrecisionTrack* enables scalable, long-duration behavioral analyses including social dynamics.

*PrecisionTrack* follows a tracking-by-detection paradigm in which subjects are localized at each timestep and associated over time. Because localization quality constrains identity assignment, jointly optimizing detection and pose estimation is critical [39]. In contrast to top-down or bottom-up pipelines that decouple these steps [5, 6], we adopt a single-stage approach inspired by *YOLO-Pose* [21], enabling pose estimation to benefit directly from advances in object detection [22-25] while reducing latency. Coupled with a modern backbone (*CSPNeXt*), this design improves receptive field and gradient flow, enhancing performance in dense scenes. In practice, we showed that this unified architecture improves both accuracy and efficiency while providing a streamlined foundation for downstream tracking and behavioral analysis.

Tracking performance is typically assessed using MOTA and IDF1 [41]. While MOTA reflects detection quality, IDF1 captures identity continuity and is more sensitive to association errors, particularly in occlusive settings. Accordingly, improvements in association predominantly affect IDF1. Our ablation analyses show modest MOTA changes but substantial IDF1 gains relative to *ByteTrack*, indicating improved identity preservation rather than detection changes. Although MOTA and IDF1 often correlate [41], our results demonstrate that optimized association mechanisms selectively enhance identity tracking. By benchmarking against *DeepLabCut, SLEAP, SORT*, and *ByteTrack*, we confirmed this and showed that *PrecisionTrack* achieves high IDF1 while maintaining competitive MOTA, supporting robust trajectory reconstruction in dense environments.

Despite these improvements, prolonged occlusions remain a key limitation, as reliably distinguishing between visually similar subjects remains an open challenge [54]. This is shown by our trajectory stitching accuracy rapidly decreasing as the number of subjects simultaneously exiting hiding spots increases. When visually similar subjects disappear simultaneously, such as when multiple individuals enter the same shelter or nesting area, our motion-based stitching algorithm alone cannot reliably disambiguate them upon reappearance. To mitigate this, we integrated the *Tailtag* system, a lightweight visual tagging approach supporting long-term identification [44], and adapted it for *PrecisionTrack* using localized detection strategies. This combination improves identity recovery and outperforms standard trackers over long durations. However, *Tailtag* is not universally applicable due to species-specific constraints. In species with distinctive features, appearance-based re-identification [54] offers a promising alternative. Future work will enable the integration of such re-identification modules into *PrecisionTrack*, leveraging its modular design.

Tracking alone is insufficient in modern ethological pipelines in which both extracting and interpreting behavior is becoming essential. We therefore introduce *MART*, a transformer-based module for real-time, subject-level behavior classification. Unlike conventional approaches (*MLP* [48], *LSTM* [47], *SimBA* [10]), which lack temporal integration or rely on fragmented predictions, *MART* models behavior as a temporal inference problem using causal self-attention [45]. Our analyses show that incorporating temporal context significantly improves performance, with an optimal window of ~30 frames (~1 s), while longer windows introduce noise likely from older/previous actions. Additionally, our results show that combining pose, visual features, and velocity improves classification relative to single-modality inputs. To further refine *MART*’s social interaction capabilities, we developed *G-MART*, a graph neural network [51]. *GMART*’s represents subjects as nodes and their relationships as edges. Through attention-based message passing, the model incorporates the context of neighboring individuals to refine interaction predictions and infer interaction partners (i.e., who interacts with whom). This introduces a relational inductive bias well suited for social behavior modeling and enables inference of both interaction presence and directionality. In practice, this provides a more expressive and integrated alternative to offline approaches.

We capitalized on this unique feature to evaluate social interactions among tracked individuals using a continuous-time movement model [53], revealing a dynamic and evolving organization of social structures within the mouse colony over time. Although our analyses were restricted to three successive 10-minute periods, they nonetheless demonstrate that social organization is not entirely stable, even over relatively short timescales. For example, while a few pairs of individuals maintained relatively consistent interactions across time (mice #1 and #23), several others exhibited marked fluctuations in interaction strength and directionality between time periods. Importantly, the interactions analyzed here primarily reflect affiliative or non-aversive social behaviors, as prolonged interactions contribute more strongly to network connectivity. Consequently, our results should not be directly interpreted in the context of classical studies of social hierarchy or dominance, which largely rely on aggressive or competitive interactions [55-57]. Instead, our approach complements these previous frameworks by enabling quantitative investigation of positive social interactions and their temporal evolution. In the future, extending these analyses to longer recordings and broader behavioral repertoires may help refine our understanding of how affiliative and competitive interactions jointly shape social organization under varying environmental conditions.

Finally, *MART* representations organize both labeled and unlabeled behaviors into structured clusters, including sub-clusters within the “Other” category, indicating that the model captures latent behavioral structure beyond predefined labels. This organization suggests that *MART* could operate in a self-supervised regime to discover behavioral patterns without manual labeling. This supports the potential of unsupervised action discovery, as illustrated by approaches such as MoSeq [11], and motivates future integration of semi-supervised or unsupervised modules to expand behavioral repertoires while preserving flexibility across experimental contexts.

In this study, we presented the development and application of *PrecisionTrack*, a unified framework for multi-animal tracking and behavioral analysis in naturalistic environments. By integrating detection, identity preservation, action recognition, and interaction modeling within a single pipeline, *PrecisionTrack* enables accurate and scalable analysis of complex social behaviors in large groups. We demonstrated its capabilities by tracking and analyzing the social dynamics of a group of 20 mice over prolonged periods, revealing structured activity patterns and interaction dynamics that would be difficult to capture using existing approaches. Overall, *PrecisionTrack* provides a comprehensive tool to quantify behavior across spatial, temporal, and social dimensions. We anticipate that its large-scale application will help refine our understanding of social organization and its impact on how individuals adapt to environmental challenges affecting health and survival.

## METHODS

### MICE datasets and benchmark

Our pose-estimation dataset consists of over 1,500 images (1536×1536 resolution) featuring 20 mice. The images are from ethological experiments conducted in the laboratory of B. Labonté at CERVO in 2022. Each localization and skeleton (**Figure 2a**) were manually labelled internally. This dataset follows Microsoft’s popular COCO formatting standard. This dataset was used to train *PrecisionTrack, DeepLabCut* and *SLEAP* before evaluating their respective performances on our benchmark. All animals’ procedures followed the Canadian Guide for the Care and Use of Laboratory Animals and were approved by the Animal Protection Committee of Université Laval.

In addition to our pose-estimation dataset, we also compiled a sequential dataset comprising 35 minutes (30/5 train/val split) of recording at 28 Hz for a total 58,800 frames (2720×2720 resolution), which correspond to approximately a million localizations, poses, and actions of socially interacting mice in an enriched open field. The labelling of mice localizations follows the MOT standard, with each instance having a bounding box, keypoints and an action annotation. The bounding boxes are defined by ρ, while the keypoints are represented by 8 (x, y) coordinates corresponding to the snout, right ear, right leg, left ear, centroid, tail base, and tail tag, respectively (**Figure 2a**). A total of five distinct actions were labelled (Grooming, Running, Interacting, Rearing and Other), we wanted to also support Fighting, but we could not find enough fighting instances in our recordings to gather an interesting dataset. This dataset was used to train and evaluate our Multi-animal Action Recognition Transformer (*MART*; **Figure 6**).

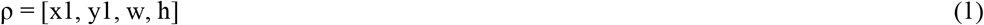

To complement these datasets, we provide a manually labelled evaluation benchmark, consisting of 20 interacting mice viewed from top-down. Here, we leveraged this benchmark to evaluate the tracking performances of *PrecisionTrack, DeepLabCut, SORT, ByteTrack* and *SLEAP* on challenging, ethological settings (**Figure 4c**). These recordings were captured during additional ethological experiments conducted in B. Labonté’s laboratory at CERVO, with a frame rate of 28Hz and a resolution of 1536×1536 pixels.

### Detection step

*PrecisionTrack* follows the Tracking-By-Detection (TBD) paradigm, composed of two main autoregressive steps: 1) Detecting instances 2) Assigning current detections to memorized trajectories [38]. As such, *PrecisionTrack*’s first step, referred to as the detection step, focusses on localizing, classifying and estimating the pose of the subjects by processing raw RBG images. Besides some pre and post processing, our detection step is end-to-end, meaning it is solely composed of optimizable parameters, maximizing *PrecisionTrack*’s generability and transferability. These optimizable parameters formed our efficient multi-task CNN architecture inspired by [25, 21].

### Neural network architecture

*PrecisionTrack*’s neural network architecture integrates recent advances in real-time object detection to balance computational efficiency and accuracy. First, it leverages the *CSPNeXt* architecture (**Figure 2d**). Introduced by [25], *CSPNeXt* is designed to maximize receptive field without significantly increasing the network’s floating-point operation (FLOP) count or its latency. As such, *CSPNeXt* relies on large convolution kernels (5×5) to expand the model’s effective receptive field, capturing broader spatial context beneficial for dense prediction tasks like object detection and pose estimation at the cost of an increase computational demand relative to narrower convolution kernels (3×3). To counterbalance this increase, it also leverages depthwise separable convolutions (**Figure 2d**). However, since depthwise separable convolutions are composed of a sequence of two (depthwise and pointwise) convolution operations, blindly switching vanilla convolution layers for depthwise separable convolution layers in any given network will effectively double its depth, potentially impeding its parallelization potential. Therefore, the backbone architecture was adjusted by moderately increasing the filter count in each layer, thus widening the network and reducing the total number of layers from 9 to 6. This configuration yields an optimal computation-accuracy trade-off by decreasing latency without compromising detection performance [25]. Lastly, to efficiently focus on the relevant features extracted by the newly widened layers, a Channel Attention mechanism is added at the end of each of them (**Figure 2d**), this has been shown to lead to an increase in performance [25].

### Multi-subjects detection and classification

*PrecisionTrack* employs an anchor-free detection mechanism, directly regressing subjects’ locations and poses from a predefined grid of priors. Compared to anchor-based approaches, which depend on predefined anchor boxes of multiple scales and aspect ratios, it enables a more direct association between detections and features extracted by the backbone [35]. Thereby enabling efficient downstream processing relying of detection’s features such as action recognition. Leveraging three distinct anchor-free heads, *PrecisionTrack* independently regresses, relative to each prior P, bounding boxes coordinates relative to each prior 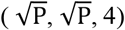, detection scores 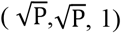 and classification scores 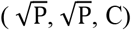, where C represents the number of possible classes. These priors correspond to unique feature maps, of dimensions (20 × 20), (40 × 40) and (80 × 80), coming directly from our three-stage feature pyramid (**Figure 3a**). By ensuring that the feature maps are symmetrical, their width (W) equals their height (H), we can express the priors as 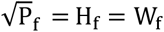. Thus, each stage of the feature pyramid processing 640×640 images produces 400, 1600, and 6400 priors, with strides of 32, 16, and 8, respectively. Resulting in a total of 8400 priors, or potential detection, per image. To mitigate the overlapping of true positive coming from this high number of potential detections, *PrecisionTrack* leverages the fast and configurable Non-Max Suppression (NMS) algorithm to filter out bounding boxes overlapping by more than 65%.

### Multi-subjects pose-estimation

Pose estimation algorithms typically follow either a top-down or bottom-up approach. Top-down (or two-stage) methods are generally more accurate, as they first detect all relevant subjects in the image and then perform single-individual pose estimation for each detection. However, the computational cost of these methods scales linearly with the number of processed individuals, leading to variable runtime. This makes top-down pose-estimator unsuitable for scenarios like ours, where the goal is to track large groups of subjects simultaneously and in real-time. In contrast, bottom-up approaches offer constant runtime, but they tend to be less accurate because of their two-stage process: 1) detect all keypoints in a single pass, 2) group them into individuals through various post-processing steps, such as Part Affinity Fields (*PAF*s) [58]. To overcome these limitations, we leverage recent advancements in both object detection [22, 25] and pose estimation [21], extending our existing multi-subject detector to support accurate pose estimation while maintaining a constant runtime and without adding additional post-processing. By adding two more anchor-free heads of shapes 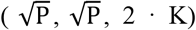 (**Figure 2c**) and 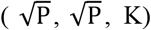 respectively, *PrecisionTrack* is able to regress keypoints directly from priors, inherently linking them to individuals. This approach thus directly resolves the keypoint-to-individual problem, eliminating the need for additional post-processing. As a result, we extend our object detector to support pose estimation while only marginally increasing the network’s computational cost at runtime.

### Subject detection, classification and pose-estimation training

During the training process, predictions for each stage of the feature pyramid are matched with ground truth through the SimOTA [22] dynamic label assignment strategy. Each decouple head has its own loss function to optimize: 1) The classification head optimizes the Quality Focal (QFL) [59] Loss (**Equation 3**), 2) the bounding box regression head optimizes the General Intersection over Union (GIoU) loss (**Equation 20**), 3) the keypoint regression head optimizes the Object Keypoint Similarity (OKS) [17] loss (**Equation 2**), 4) both the detection and keypoints confidence regression heads optimize the Binary Cross Entropy (BCE) loss. As such, the final loss function is composed of a weighted sum of each decoupled head’s loss, as defined by **Equation 4**.

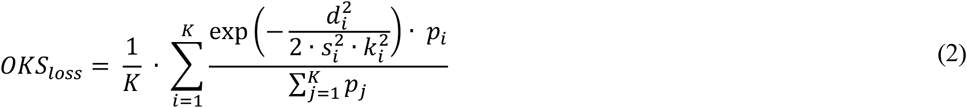

Where:

- K : Total number of keypoints.
- *d*_*i*_ = ‖*gt*_*i*_ − *pr*_*i*_‖_2_
- *k*_*i*_: Weight of the i-th keypoint.
- *p*_*i*_: 1 if the i-th ground truth is present, else 0.

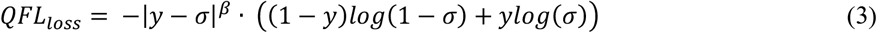

Where:

- σ: The model’s prediction.
- y: The ground truth.
- β: Down-weighting rate’s hyperparameter (set to 2).

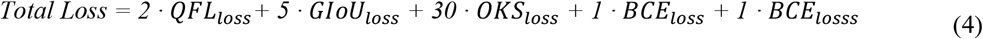

Our two-stage training process closely follows [22]’s. Its first stage uses strong data augmentations, Mosaic and MixUp (**Figure 2b**), to teach the model that relevant subjects could be anywhere and at any scale. This step is crucial for minimizing common biases found in human-taken images, where objects of interest are often centered, non-occluded, and fully in the frame. It has also been shown to minimize the risk of learning these biases rather than the actual discriminative features necessary to accurately localize relevant subjects in real-world conditions [25]. The second training stage leverages weaker geometric augmentations, such as cropping and flipping, fine-tuning the model for the specific task at hand.

We adopt a hyperparameter-free cosine learning rate schedule, preceded by a brief warm-up period. Given our limited computational resources, a single RTX 3090, we utilized the linear learning rate scaling approach proposed by [60]. This allowed us to effectively replicate the performance benchmarks of *YOLOX* [22] on the COCO dataset [19] while using a fraction of the computational power.

### Detection and pose-estimation evaluation

An optimal multi-animal pose tracking algorithm’s detection step should prioritize three core qualities: 1) high detection coverage, meaning detecting as many subjects as possible in each frame, accurate pose-estimation on each of the detected subjects, and 3) low latency to support real-time tracking. While the standard COCO metrics [19] technically encompass these qualities, they can be less intuitive to understand compared to other widely used metrics. Therefore, we report additional, flexible metrics with greater interpretability.

Specifically, we evaluated our model based on a weighted sum between the detection’s F1 score and the detected subject’s Object Keypoint Similarity (OKS) metric (**Equation 13**). To closely reflect the inference scenario in our testing pipeline, our predicted detections are matched to labels using *PrecisionTrack*’s IoU-based cost function (**Equation 14**) and the Hungarian Algorithm [61].

For pose-estimation accuracy, we prioritize OKS and PCK over RMSE (**Figure 2e**), commonly used in other methods [6]. As both OKS and PCK are normalized by subject size, reducing the impact of regression errors on larger subjects.

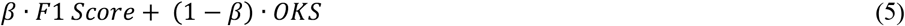

Where:

- *β* = 0.5

### Feature extraction

As explained in the Multi-subjects detection and classification section, the backbone outputs 8,400 feature vectors, one per prior. Separate heads directly regress these vectors to predict each subject’s class, pose, and location (**Figure 3a**). Therefore, these vectors necessarily encode semantics allowing downstream tasks to accurately locate, classify, and estimate poses. Consequently, these features could be interpreted as processed representations of each subject’s appearance and offer rich inputs for other downstream tasks such as action recognition, enabling an efficient analysis pipeline.

An unintuitive finding has been that our one-stage, anchor-free and FPN-based object detector encoded uncoherent feature maps with respect to the inferred classes, localizations and poses. Specifically, features associated with highly overlapping bounding boxes, e.g., redundant localizations of the same entity, remained distant in the feature space (**Figure 3b**). This mismatch between detection and latent-space representation means that features linked to the same entity can drastically vary across timesteps, inevitably impacting the performances of sequential downstream tasks.

To solve this incoherence problem, we introduce a new convolution-based head trained to cluster relevant features by optimizing a triplet loss with easy and semi-hard mining. During inference, the resulting clustered features are normalized and aggregated via a weighted pooling mechanism taking into consideration each feature’s associated confidence score. This process stabilizes each timestep’s features, producing normalized and coherent feature maps (**Figure 3b**). Because the newly introduced head’s clustering objective does not directly align with the network’s other head’s objectives, e.g., detection, classification and pose-estimation, we propose to replace the standard multi-task JDE approach popularized by [34, 35] by a curriculum-based strategy. By first training the network as previously described in the *Multi-subjects detection and classification and Multi-subjects pose-estimation* sections, and then training the clustering head separately, on frozen embeddings, *PrecisionTrack* enhances and stabilizes its feature extraction ability without impacting its accuracy.

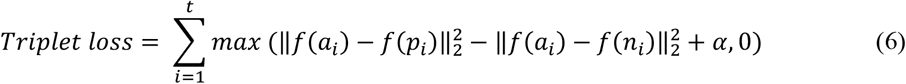

Where:

- *a*: The anchor, the reference point.
- *p*: The potitive, a point with the same identity as the anchor.
- *n*: The negative, a point with a different identity than the anchor.
- *α* = 0.2

### Association step

The association phase, which follows detection in the TBD pipeline, aims at pairing new detections to their respective live trajectories. Although conceptually simple, the task becomes combinatorially challenging whenever the detection count diverges from the number of active tracks. Following a recent trend in the Multi-Object Tracking (MOT) field [40], *PrecisionTrack*’s association step prioritizes a high recall approach. Therefore, while previous multi-animal pose tracking algorithms opted for systematically filtering out low-confidence detections before the association process [5, 6], *PrecisionTrack* salvages them in a second association step designed to perform low-confidence assignation. This method has been shown to yield more robust tracking performances in scenarios where the visible entities are partially occluded, such as within crowds or in complex environments [40].

### The projection substep

Most current state-of-the-art TBD algorithms split the association step into two substeps: the projection substep and the matching substep. In the projection substep, motion estimation models are leveraged to project the tracked trajectories in the current frame, facilitating eventual matching [38]. Among these projection models, the Kalman Filter [42] remains the preferred choice due to its efficiency, reliability, and low computational cost. Especially in the multi-object tracking (MOT) community, whose benchmarks mainly include pedestrians and vehicles. The Kalman Filter’s assumptions of linear motion and constant velocity align well with the typical movement patterns of these objects, such as pedestrians walking along sidewalks or vehicles following lanes.

However, these assumptions do not hold as well in multi-animal pose tracking where tracked subjects are often freely roaming in an open field while being recorded from a top-down view (**Figure 4b**). This could lead to sudden changes in the subject’s direction and velocities (**Figure 4b**). As such, we argue that this significant shift in data dynamics imposes a corresponding shift in the leveraged motion projection techniques. To bridge this domain gap, we propose a simple, yet effective motion algorithm called the Dynamic Kalman Filter (DKF), which incorporates 2D Newton’s laws of motion into the vanilla Kalman filter to more accurately project trajectory dynamics over time, even in cases of abrupt 2D changes in both direction and velocity (**Figure 4b**).

The DKF updates each trajectory’s dynamics after each association step using the following:

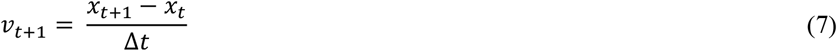

Where:

- *v*: The trajectory velocity
- *x*: The trajectory position

For numerical stability, given the frequent directional changes observed in animals, an exponential moving average (EMA) is then applied to the velocity estimate:

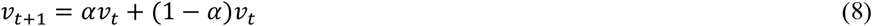

This smoothed velocity estimate is then used to infer the trajectory’s accelerations using Newton’s second law of motion:

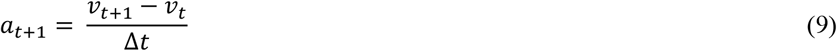

Where:

- *a*: The trajectory acceleration

To further accommodate the complexities of animal tracking, we also adjusted the Kalman Filter’s noise estimation. We set the initial state of the covariance matrix P to 1000 times the identity matrix, representing high uncertainty in the absence of prior knowledge about the trajectory’s states. P is updated at every time step using **Equation 12**. The noise covariance Q is scaled by 0.1 and dynamically adjusted based on the time step Δt. This models the position noise growing quadratically and the velocity noise linearly in respect to eventual delays between measurements, allowing us to account for potential broken trajectories. The measurement H and measurement noise covariance R are both set to diagonal matrices with values of 1.0, assuming equal noise in both the x- and y-axes.

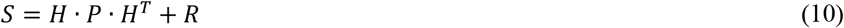

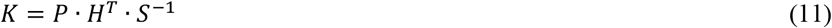

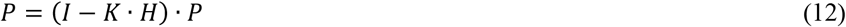

### The matching substep

The matching substep involves linking the projected trajectories to new detections. Typically, matching costs are defined by the Intersection over Union (IoU) (**Equation 18**) between the bounding boxes of detections. However, we argue that relying solely on bounding boxes overlooks critical information that could lead to tracking errors. For instance, two trajectories may overlap and exhibit similar IoU scores with newly acquired detections; however, the two detected subjects might not even be facing the same direction. As such, incorporating additional information, such as the pose of the subjects, can enhance the robustness of the association step. Thus, we propose to complement the standard IoU scores with our slightly modified version of the Object Keypoint Similarity (OKS) loss function [21]. Our implementation of OKS normalizes the Euclidean distances (**Equation 13**) by the trajectory’s areas *A*. The exponential term in the OKS calculation is also multiplied by the weight of each detection *w*.

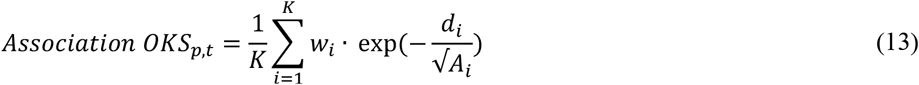

Where:

- *d*_*i*_ = ‖*p*_*i*_ − *t*_*i*_‖_2_
- *A*_*i*_ = *h*_*i*_ · *w*_*i*_

By combining a weighted sum of the IoU and OKS metrics with a lambda of 0.5, we achieved a richer matching cost while retaining the advantages of the IoU metric, such as the ability to easily identify non-overlapping trajectory-detection pairs.

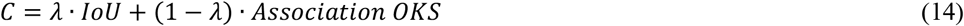

### Trajectory stitching

Trajectory stitching is an optional plugin for *PrecisionTrack* tracking pipeline. It allows the reconstruction of sparse trajectories for classes of subjects confined within the camera’s field of view. For example, if a user wishes to track *n* marmosets (*Callithrix jacchus*) in their home-cage while also counting the number of new researchers entering the camera’s field of view, they can configure the stitching plugin to only consider the *n* marmosets. This ensures that whenever a new marmoset is detected, e.g., emerging from a hideout, *PrecisionTrack* will link it to a loss marmoset trajectory instead of considering it as a new trajectory, preventing an incorrect increment of the marmoset count.

While offline methods such as *DeepLabCut*, by processing whole recordings as once, can leverage clever techniques such as network flow minimization algorithm to solve this problem, online methods like *PrecisionTrack*, since it does not have access to global trajectories, must instead solely rely on trajectory’s past information to eventually stitch them on the fly.

Working within this constraint, we developed a fast and flexible heuristic algorithm called Search-Based Stitching (SBS) that delivers strong practical performance. SBS defines a dynamically evolving search zone around lost trajectories’ last known location. These search zones expand over time according to the lost trajectories last recorded dynamics (**Figure 5a**). To minimize the risk of excessively expanding the search zone, which could lead to a higher rate of false-positive matches, the rate of expansion decays at an exponential rate (**Equation 15**).

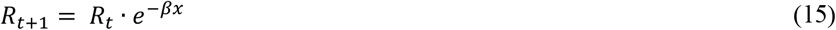

Where:

- *R*: The search zone expansion
- *β*: Decay parameter, default to 0.1

Additionally, the search zones expand asymmetrically, guided by each side’s cosine similarity with the lost trajectory’s last recorded EMA 2D velocities (**Equation 16**).

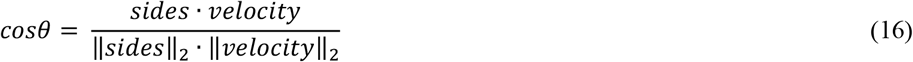

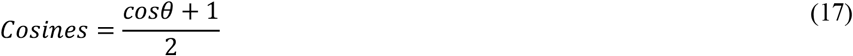

This biases the search towards the area the lost tracklet is most likely heading for (**Figure 5a**). Lastly, to account for the frequent use cases where multiple search zones are active and/or multiple new trajectories are detected at the same time, we cast the search zone-to-new trajectory linking as an assignment problem, which we solve using the Hungarian algorithm [61]. The cost matrix for the association is based on our implementation of a normalized Generalized Intersection over Union (GIoU) (**Equation 19**) between each active search zone and each new relevant detection.

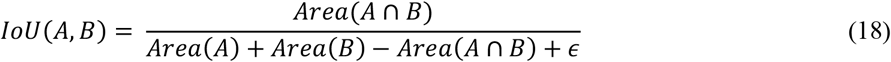

Where:

- 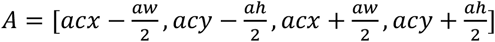
- 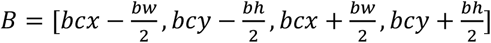

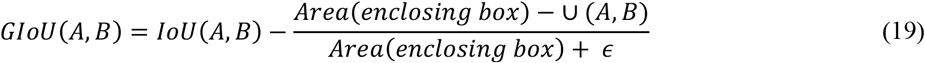

Where:

- *enclosing box* = min(*A*_1,_ *B*_1)_, max (*A*_2_, *B*_2_)

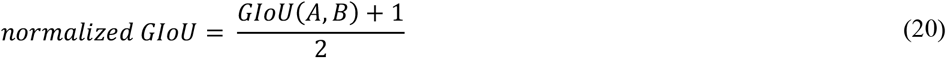

### *Tailtag* re-identification

While the *Tailtag* system is shown to be effective and non-behavioral altering [44], it still has some weaknesses. Mainly; 1) the detection process, while robust, is not impervious to mistakes, and 2) searching for tags on the whole frame is computationally expensive. In fact, leveraging OpenCV’s ArUco detection functionalities as is would bottleneck *PrecisionTrack*’s latency and harm its ability to run at real-time speed. Therefore, we opted to refine [44]’s approach in two ways. First, leveraging the fact that we are already tracking the tags’ locations via our pose-estimation pipeline (**Figure 2a**), we were able to significantly reduce the marker search area from the whole frame to only regions around the subject’s tags (**Figure 5b**). Additionally, we built a timeout priority queue, ensuring that the most uncertain trajectories have re-identification priority while also ensuring that the *Tailtag* re-identification will not bottleneck the whole system’s latency via a timeout mechanism. Second, the system perpetually accumulates evidence in the form of detections on each of the subject’s tag areas. Tags are assigned to subjects only when confidence thresholds are met. This assignment could lead to what we define as a correction, meaning that a subject (given enough accumulated evidence) could get assigned to another one’s tag. In this scenario, the *Tailtag* re-identification system declares a correction and the subject’s identities are corrected accordingly. Paired with the *Tailtag* re-identification plugin, *PrecisionTrack* maintains reliable tracking performances over extended periods (**Figure 5d**), while remaining computationally efficient enough to remain real-time (**Figure 4c**).

### Tracking evaluation

Tracking performances are assessed using the CLEAR metrics [62], which have been the standard for multi-object tracking (MOT) evaluations for over a decade. Specifically, we employ the Multiple Object Tracking Accuracy (MOTA) metric (**Equation 30**), which quantifies three main types of errors that can occur during tracking:

1. False Positives (*FP*): Instances where the algorithm incorrectly tracks irrelevant objects.
2. False Negatives (*FN*): Instances where the algorithm fails to track relevant objects.
3. Identity Switches (*IDSW*): Instances where the algorithm misidentifies two trajectories, resulting in swapped identities.

As such, MOTA focuses on assessing the overall quality of the tracking algorithm:

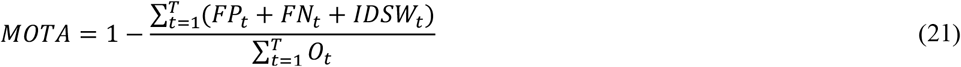

Where:

- *FP*_*t*_: Total number of false positives at time t.
- *FN*_*t*_: Total number of false negatives at time t.
- *IDSW*_*t*_: Total number of identity switches at time t.
- *O*_*t*_: Total number of visible objects at time t.
- *T*: Total number frames.

Our evaluation pipeline also incorporates the IDF1 score (**Equation 31**), which focuses on evaluating the algorithm’s ability to maintain the correct identities of tracked objects over time:

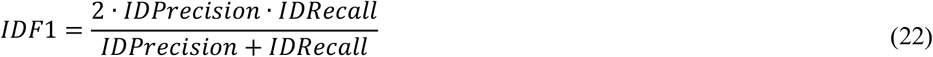

### Training Pipeline

A typical *PrecisionTrack* training workflow requires three key inputs: 1) Users define their experiment’s metadata (e.g. their subject’s species, actions, keypoints and skeleton). 2) Users label their data using popular and established computer vision labelling suites such as CVAT or LabelStudio. 3) Users register their COCO formatted training dataset as well as their metadata file into the PrecisonTrack’s settings. With these inputs, users can employ the provided training and testing tools to train, validate and test their *PrecisionTrack* system. Once satisfied with the performance, users can use the deploy tool to convert the PyTorch checkpoint into a fully optimized *PrecisionTrack* engine. The deployed system can then be used to automatically perform inference on new, unseen recordings.

### Active Learning

Users are encouraged to train their own customized *PrecisionTrack* system iteratively. The process begins with the annotation of 50–100 frames randomly sampled across the recordings. Importantly, consecutive frames should be avoided, as they are often redundantly similar and thus provide little to no value during the optimization process. Once this initial labeled set is complete, a first version of *PrecisionTrack* can be trained to support subsequent labeling. Users can then apply this model to new recordings, identify frames with uncertain detections (e.g., predictions with 50–70% confidence), and refine only those. The corrected predictions are incorporated into the dataset as additional labels. This iterative procedure, known as *active learning*, can be repeated as often as necessary. It offers two main advantages: 1) It ensures that users label only informative and challenging examples, which maximizes model improvement while avoiding redundant data. 2) It accelerates the labeling process by presenting users with predictions to refine rather than requiring annotation from scratch.

### Action recognition

The Multi-animal Action Recognition Transformer (*MART*) is designed to model the inherently dynamic structure of animal actions. Modern transformer-based natural-language-processing methods achieve state-of-the-art performance by selectively reasoning over linear sequences [45, 46]. By repurposing transformer decoder blocks to operate on subjects’ state representations rather than tokenised words, *MART* captures the temporal dependencies that characterise behavior. Whereas traditional action-recognition benchmarks focus on sparse predictions for a single actor [63, 64], *MART* jointly predicts actions for large groups of individuals at frame-level temporal granularity. This shift introduces three challenges: (1) the inability to isolate specific actions yields severe class imbalance, since some behaviors naturally occur far more often than others; (2) the model must scale jointly with context length and number of subjects; and (3) restricting the input to subjects’ poses provides limited context, omitting appearance and social cues required to disambiguate actions in group settings.

We address the first challenge by grouping rare or undesired actions into a single “Other” class and by enforcing class uniformity at the sequence level. During training, each sample is constructed by first drawing an action *a* with probability *p*(*a*) ∝ 1/*n*_*a*_ (or uniformly *p*(*a*) = 1/∣ 𝒜 ∣ when no weighting is applied), and then sampling a sequence of length *T* centred on a frame in which *a* was annotated. This procedure mitigates the over-confidence typically observed when training on imbalanced data. Across our hyper-parameter sweep, the best accuracy–latency trade-off was obtained with four decoder blocks, four attention heads per block, and a context window of *T* = 30 frames (**Figure 6b**). Because no pretrained transformer is available at this scale of multi-animal behavior, we trained the network from scratch and obtained strong performance (**Figure 6c**) without the next-token-prediction pretraining phase that is standard for language models.

We address the second challenge by parallelising across the subject dimension. For a batch of *B* scenes containing *N* subjects each, the per-subject state sequences of length *T* are reshaped to (*B* · *N, T, d*) before entering the decoder, exposing subjects as independent items in the leading batch dimension. Because transformer blocks are heavily optimised for GPU execution, this trivially leverages tensor cores and yields throughput that scales near-linearly with *N*. The only non-parallelisable component is the depth of the decoder stack, which we therefore limited to four blocks, achieving an average per-frame latency of 0.0117 s on an RTX-3090.

We address the third challenge by exploiting our feature-extraction pipeline to ablate alternative state-embedding compositions. For each subject *i* at time *t*, the state comprises three modalities: (1) pose key-points 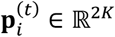, (2) appearance features 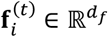 extracted from the tracking backbone, (3) instantaneous velocities 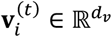. Each projected through a modality-specific encoder *ϕ*_·_ composed of a linear projection, a position-wise MLP, and layer normalisation. The three projections are concatenated and linearly mixed before the addition of a learnable positional embedding **u**^(*t*)^:

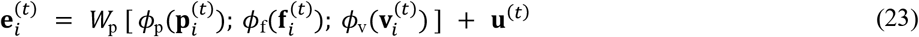

Where:

- [·; ·] denotes concatenation along the feature dimension.

### Graph-based refinement of social interactions

While *MART* recognises the actions of individual subjects from their per-frame state sequences, it does not encode the *who-with-whom* of a social interaction: a frame in which subject *i* is labelled “Interacting” carries no explicit information about the partner subject *j*, nor about who initiated the contact. Inferring this directional structure requires reasoning jointly over all subjects in the scene. To this end, we introduce *G-MART* (Graph-*MART*), a graph-based head that consumes *MART*’s per-subject embeddings at the last time-step of the context window and produces (i) a refined per-subject social-action distribution and (ii) a directed interaction graph over the *N* subjects.

Social interactions in mice are mediated by user-defined spatial-priors (snout-to-snout, snout-to-ears, centroid-to-centroid, etc.), which provide a strong inductive bias for the relevance of any pair of subjects. For each ordered pair (*i, j*) and each user-defined keypoint pair *k* = (*k*_src_, *k*_tgt_), *k* = 1, …, *K*, we compute a body-scale-normalised distance

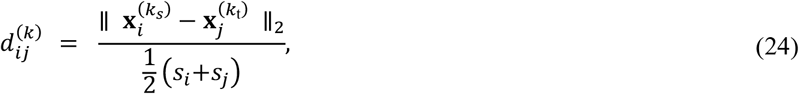

where 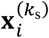 is the image-plane location of keypoint *k*_s_ on subject *i, s*_*i*_ is the subject’s bounding-box scale. When either keypoint is occluded, the visibility gate

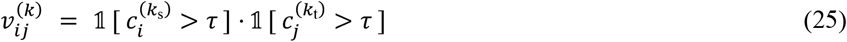

suppresses the corresponding distance, 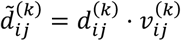, where 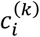 is the keypoint detection confidence and *τ* is a fixed threshold. The centroid–centroid distance is exposed as a separate, ungated *distance prior δ*_*ij*_, which guarantees that every pair receives at least one informative spatial signal even when all keypoints are occluded.

The full vector of per-pair spatial priors 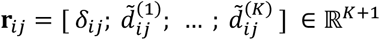 is mapped to a relationship weight 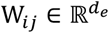 by a two-layer MLP with GELU activation and layer normalisation, W_*ij*_ = MLP(**r**_*ij*_). Relationship weights are normalized by linearly projecting W_*ij*_ and applying a per-row softmin so that geometrically closer (more relevant) partners receive larger weights:

Let 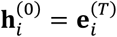 denote the *MART* embedding of subject *i* at the final time-step of the context window, and 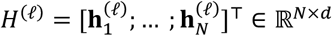. The graph encoder is a stack of *L* non-causal transformer blocks operating over the *N*-subject sequence, in which the standard self-attention is biased by the relationship weights *W*_*ij*_:

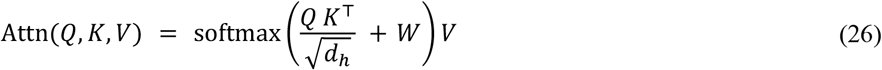

*G-MART* is trained end-to-end with a weighted sum of two focal losses [52]. The node loss is a multi-class focal loss over labelled subjects 𝒱_lab_ with optional per-class weights ***α***^cls^ and label smoothing *η*; the edge loss is a binary focal loss over the set of valid pairs ℰ_val_ = {(*i, j*): *i* ≠ *j, i, j* valid}:

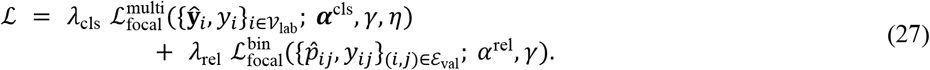

We use focusing parameter *γ* = 2 throughout, an edge-class weight *α*^rel^ = 0.95 to counter the strong negative skew of the interaction graph (most pairs are non-interacting), and *λ*_rel_ ≥ 1. During *G-MART* training the *MART* backbone is frozen, so that *G-MART* learns purely a relational refinement on top of *MART*’s temporal representations, preserving *MART*’s per-subject action distribution while adding directed-interaction inference at negligible additional cost.

### Ecological study

The continuous time movement model was developed based on the annotated *PrecisionTrack* data where interacting individuals were identified. From a modelling perspective, we adapted Yang et al. [53], to quantify the interactions an individual has on another. Specifically, for each 10-minute blocks we measured the number (*n*) and length (*l*) of the period an individual was interacting with another and weighted as

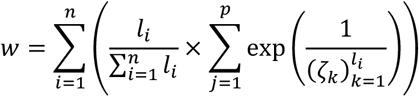

where 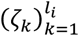 is a sequence of time steps going from 1 to *l*_*i*_. In words, *w* weights the importance of an individual has on another one by accounting for the number of periods they are together and how long these period last. In this calculation, two individuals that are together for a long period will be given more importance than if they are together for the same amount of time but separated in multiple short periods.

### Computational Resource Usage

All experiments were conducted on a machine equipped with 128 GB of RAM, an Intel i7-10700k CPU, and an RTX-3090 GPU. The *PrecisionTrack* algorithm requires approximately 4 GB of RAM and 2.5 GB of VRAM for execution. Additionally, saving the tracking results incurs a disk space cost of about 7.3 MB per minute when monitoring 20 subjects.

## Code availability

*PrecisionTrack* is fully open-sourced: VincentCoulombe/precision_track.

Our repository includes:

- Detailed installation and usage instructions.
- Pre-configured Google Colab notebooks.
- Tutorials and documentation for training, testing, deploying and tracking.

The repository is actively maintained and integrates automated testing, formatting, and continuous integration workflows.

## Data availability

Datasets, models and benchmarks are openly available. Specifically:

- The MICE dataset: [MICE dataset - Google Drive](https://drive.google.com/drive/folders/18Ikogjyeo_CRe9Z_iQfqYrfOfGt_dMma).
- The *Tailtag system* plans: [Checkpoints - Google Drive](https://drive.google.com/drive/folders/1fpKgfnE3xD9xicfxzWdXDmA1p5lE8qmm).
- Pretrained models and training checkpoints: *Tailtag*s - Google Drive.

All resources can also be accessed via the *Resources* section of the *PrecisionTrack* repository.

## Acknowledgements

BL holds a Sentinelle Nord Research Chair, is supported by the Canadian Institutes of Health Research (Grant No. PJT-451728 and PJT-451858), and the Natural Science and Engineering Research Council of Canada (Grant No. RGPIN-2019-06496) and receives Fonds de Recherche en Santé du Québec (FRQS) Senior salary support. BG holds the Canada Research Chair in S*MART* Biomedical Microsystem and is supported by the Natural Science and Engineering Research Council of Canada (Grant No. RGPIN-2022-03984). This work is made possible by the MMLab’s open-source MMEngine [65], MMCV [66], MMdetection [67], MMdeploy [68] and MMPose [69] toolkits. We also thank Arturo Marroquin Rivera for his comments on the manuscript and Maggy Shatov and David-Alexandre Roussel for refining the MICE dataset’s annotations and the data-driven ArUco reading system, respectively.

## Author contributions

B.L. supervised the project and wrote the manuscript. BG supervised the project and reviewed the manuscript. GB developed the continuous time movement model. VC developed *PrecisionTrack, MART* and *G-MART*, performed the analyses, made the figures and wrote the manuscript. MSM labelled the MICE dataset. KA labelled the MICE dataset. QL and MRP contributed to the animal experiments and video data generation. All authors contributed to the preparation of the manuscript.

## Disclosures

The authors declare no competing financial interests.

## Notes

Disclosure Statement: Authors declare no conflict of interests.

### Competing Interest Statement

The authors have declared no competing interest.

### Summary of Updates

This version of the manuscript has been revised to update the following: - Additional social dynamic analysis results - New comparisons (SimBA, ByteTrack, SORT) - G-MART

